# MAGNETO: an automated workflow for genome-resolved metagenomics

**DOI:** 10.1101/2022.05.06.490992

**Authors:** Benjamin Churcheward, Maxime Millet, Audrey Bihouée, Guillaume Fertin, Samuel Chaffron

## Abstract

Metagenome-Assembled Genomes (MAGs) represent individual genomes recovered from metagenomic data. MAGs are extremely useful to analyse uncultured microbial genomic diversity, as well as to characterize associated functional and metabolic potential in natural environments. Recent computational developments have considerably improved MAGs reconstruction but also emphasized several limitations, such as the non-binning of sequence regions with repetitions or distinct nucleotidic composition. Different assembly and binning strategies are often used, however, it still remains unclear which assembly strategy in combination with which binning approach, offers the best performance for MAGs recovery. Several workflows have been proposed in order to reconstruct MAGs, but users are usually limited to single-metagenome assembly or need to manually define sets of metagenomes to co-assemble prior to genome binning. Here, we present MAGNETO, an automated workflow dedicated to MAGs reconstruction, which includes a fully-automated co-assembly step informed by optimal clustering of metagenomic distances, and implements complementary genome binning strategies, for improving MAGs recovery. MAGNETO is implemented as a Snakemake workflow and is available at: https://gitlab.univ-nantes.fr/bird_pipeline_registry/magneto.

**IMPORTANCE:** Genome-resolved metagenomics has led to the discovery of previously untapped biodiversity within the microbial world. As the development of computational methods for the recovery of genomes from metagenomes continues, existing strategies need to be evaluated and compared to eventually lead to standardized computational workflows. In this study, we compared commonly used assembly and binning strategies and assessed their performance using both simulated and real metagenomic datasets. We propose a novel approach to automate co-assembly, avoiding the requirement for *a priori* knowledge to combine metagenomic information. The comparison against a previous co-assembly approach demonstrates a strong impact of this step on genome binning results, but also the benefits of informing co-assembly for improving the quality of recovered genomes. MAGNETO integrates complementary assembly-binning strategies to optimize genome reconstruction and provides a complete reads-to-genomes workflow for the growing microbiome research community.

## INTRODUCTION

Genomes are a valuable resource for characterizing and understanding the diversity, ecology and evolution of microbial organisms in the laboratory as well as in natural environments. As culture-based approaches have been historically used to recover genomes and enrich reference databases, current knowledge from most reference bacterial genomes come from axenic cultures. However, despite the improvement of culture-based approaches to cultivate novel microorganisms, the number of organisms that can be isolated and cultivated remain mainly constrained by specific growth conditions. Depending on the considered environment, it is estimated that a proportion of only 0.1 to 1% of all microbial genomes could be cultivated (1, 2).

The rise of metagenomic studies, thanks to the rapid development of high-throughput shotgun sequencing, has allowed direct access to the diversity and functional potential of naturally occurring microorganisms, bypassing the cultivation bottleneck. For more than a decade, various studies have reconstructed genomes from metagenomes and contributed to describe thousands of novel microbial clades belonging to diverse environments, such as in the human gut (3), in soils and in aquatic environments (4, 5).

The reconstruction of these draft genomes, commonly called Metagenome-Assembled Genomes (MAGs), has now become a common approach, with numerous software developed during the last decade (6, 7, 8, 9). As for the reconstruction of genomes from single organisms, MAG reconstruction can be split into two main steps: first, the *assembly* of the reads obtained from the sequencing into longer sequences called *contigs*; second, the *binning* of these contigs into MAGs, mainly using their compositional and/or abundance similarities. However, MAGs reconstruction can face several limitations including gaps, sequencing errors, local assembly errors, contigs chimeras and bin contamination, i.e. the inclusion of contigs belonging to different genomes in the same bin. The binning of contigs may also miss genomic regions in which nucleotidic composition differs significantly from the genome average, such as ribosomic RNA regions, or mobile elements (10). These limitations can be partially addressed by several quality checkpoints, misassemblies detection, and manual curation (11).

In addition, low abundance organisms are usually harder to recover, due to limited reads information during the assembly process (12). When shallower sequencing is performed (i.e. the predefined number of bases the sequencer will output is low), reads from low-abundant genomes will be rare, and thus their assembly into contigs will be more difficult, as assemblers tend to consider these reads as erroneous and discard them. A common approach to increase the abundance of rare reads is to adapt the assembly strategy, that is not assembling a unique metagenomic sample (single-assembly), but *co-assembling* several samples together. Co-assembly will then tend to increase the number of occurrences of rare reads, and consequently incorporate them into resulting contigs, thereby capturing a higher fraction of the diversity within the samples. Co-assembly strategies have been instrumental for recovering higher numbers of MAGs (13, 14); however, this approach increases the probability to generate fragmented assemblies (15, 12).

The genome *binning* process consists in classifying contigs usually based on similarities of their sequence composition, their abundance, or their taxonomic affliation. In most existing softwares, binning is performed using two main metrics, namely sequence composition (6) and contigs abundance (7). Sequence composition is defined as the frequencies of all 4-nucleotides within the contig sequence, called TNF (for TetraNucleotide Frequency). The abundance (or co-abundance) represents the mean vertical coverage of the contig in one (or several) sample(s). Other metrics, such as taxonomic affliation of the contigs, may also be used to determine which contigs belong to the same bin (16). The principal differences between existing binning softwares usually rely on the algorithm used to group contigs into genome bins. Most successful software have used density-based clustering (17), Gaussian mixture models (7), affinity propagation (18), or graph clustering (9). Other methods can also perform binning on genes rather than contigs, relying on the presence of co-abundant genes within metagenomes, such as canopy clustering (19) and MSPminer (20). The objects reconstructed by these methods may not be qualified as MAGs, and are commonly referred as Metagenomic Species Genomes (MGS) and Co-Abundance gene Groups (CAGs).

Extracting knowledge from raw metagenomics data requires to handle several specific tasks, from assembly to gene calling and annotation, each of them often performed using dedicated software. Today, dedicated workflows for these tasks start to emerge (21) but they are still not widely adopted by the community. Users commonly face choosing, configuring, and running different tools, which can be challenging and time-consuming. Recently, several metagenomics workflows have been developed (22, 23, 24, 25), often using specific default parameters for each integrated software. However, these workflows usually suffer from limits towards either the assembly step or the genome binning step. In workflows allowing to perform co-assembly, sets of samples to co-assemble have to be determined and manually specified by the user, implying some *a priori* knowledge about the microbial ecosystem under study. Besides, only a single workflow (25) allows the user to compute co-abundances from metagenomes that have not been included in the assembly. As computing coabundance profiles of contigs from multiple metagenomes may increase the precision of the metric (9, 26), the impossibility to compute large-scale co-abundance may be considered a limitation of these workflows.

Here, we present MAGNETO, a fully automated workflow for genome-resolved metagenomics, implementing a co-assembly module that integrates a non-supervised method to define sets of samples to co-assemble without *a priori* knowledge. It also includes complementary strategies to compute abundance metrics from one to n metagenomes, even if they do not participate in the assembly process. In this study, we tested our co-assembly module on a set of marine metagenomes, against a co-assembly relying on existing knowledge. We also benchmarked four different assembly-binning strategies for MAGs reconstruction, on diverse datasets ranging in complexity, from a mock dataset representing a small bacterial community to human gut microbiome communities.

## RESULTS

### Determining co-assemblies using metagenomic distances

In (13), the authors studied the abundance of diazotrophic bacteria in oceanic surface metagenomes and showed that nitrogen fixation is an important feature of the prokaryotic communities living in ocean surface. As microbial genetic distances often co-vary with geographic distances in several habitats (27), co-assemblies were performed based on the geographic coordinates of the metagenomes, i.e. metagenomes belonging to the same oceanic region were co-assembled. In the euphotic zone, an average higher microbial community similarity within than across ocean regions has been observed at global scale, although a separation by regional origin is unclear (28) as other environmental factors (e.g. ocean currents) can modulate genetic proximity between populations (29). In consequence, two metagenomes geographically close do not necessarily share the highest proportion of genomes, and two metagenomes belonging to the same ocean region may not be closer to metagenomes from other regions.

Given that the main goal of co-assembly is to increase the proportion of reads belonging to a given strain or species, we propose to identify sets of samples to coassemble using metagenomic distances. To the best of our knowledge, very few studies have used sequence-based compositional distances to guide metagenomic co-assembly. Historically, metagenomic compositional distances have mainly been used to compare metagenomic samples (30) or MAGs (14), but not to actually guide the co-assembly process. Though, a few recent studies have started to use metagenomic-based distances combined with clustering to guide the co-assembly process of metagenomes (31, 32, 33), while another study has used metagenomic distances to guide the co-binning (or co-mapping) process (34). Here, we computed distances between metagenomes using Simka (35), and identified optimal clustering solutions using the Silhouette index (36) to delineate unsupervised sets of samples to co-assemble. Applying this approach on the same set of metagenomes (n=93) as in (13), we identified 24 optimal clusters. This number of clusters is significantly higher compared to the 12 clusters (Fig. 1A) based on the oceanic regions, which suggests that a different partition may be more relevant for co-assembly. As this optimal clustering generates smaller clusters, in order to insure a fair comparison between both approaches, we further identified a sub-optimal clustering (supp.fig S1) whose number of co-assembly sets is comparable to the number of oceanic regions used in (13). This second clustering identified 11 clusters, which did not match the oceanic regions previously defined (Fig. 1A).

**FIG 1.**
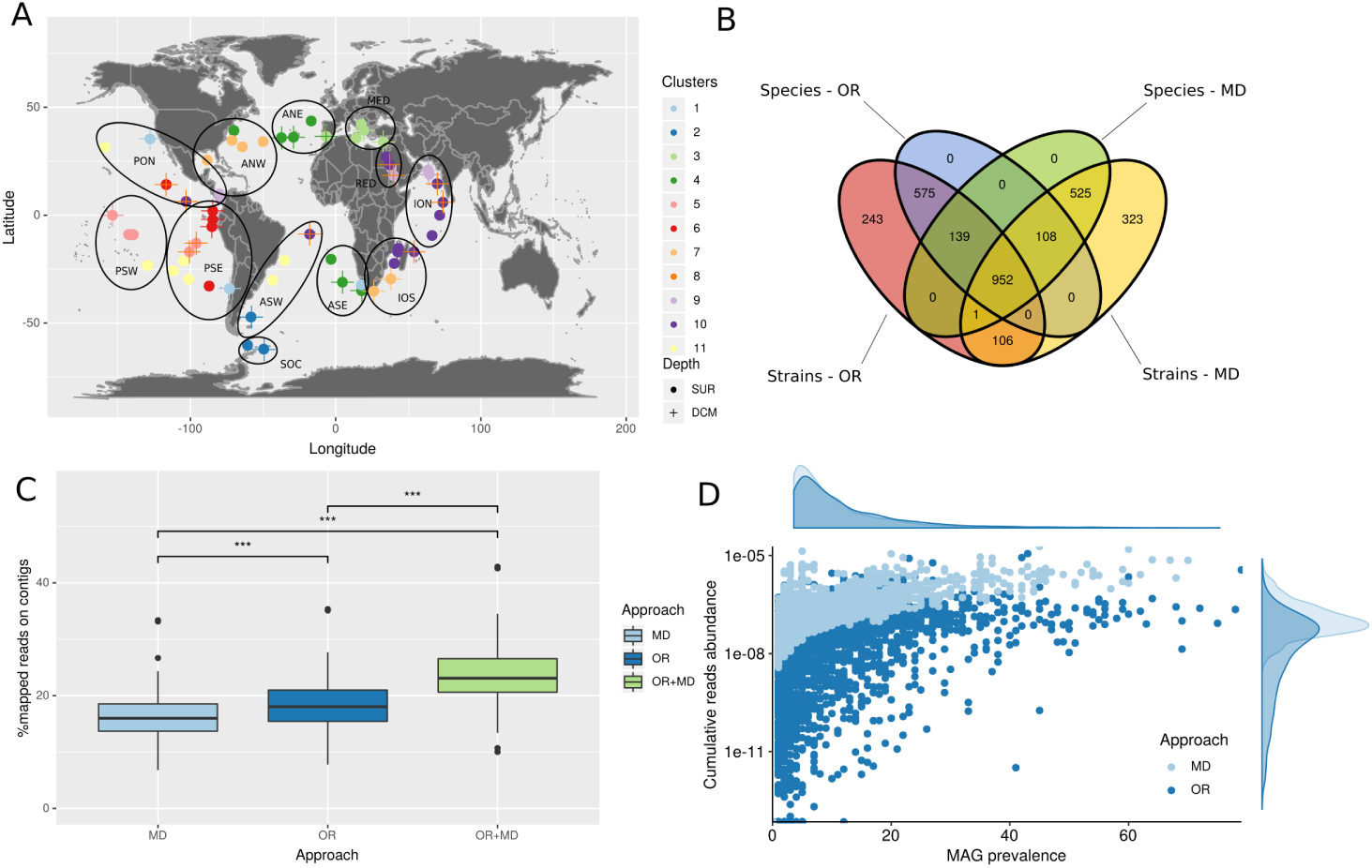
Evaluating the metagenomic distance-based (MD) approach against the Oceanic Regions (OR) approach for delineating groups of samples to co-assemble. **(A)** Repartition of clusters obtained with Simka. Each dot represents a metagenome obtained at a sampling station, with metagenomes located at Surface (SUR) represented as dots, and metagenomes situated at the Deep Chlorophyll Maxima (DCM) depth as crosses. Colours represent the cluster to which the metagenome belongs to. Oceanic Regions are represented as dark circles: ANE = Atlantic North-East, ANW = Atlantic North-West, ASE = Atlantic South-East, ASW = Atlantic South-West, ION = Indian Ocean North, IOS = Indian Ocean South, MED = MEDiterranean Sea, PON = Pacific Ocean North, PSE = Pacific South-East, PSW = Pacific SouthWest, RED = RED sea, SOC = Southern OCean. **(B)** Repartition of the common MAGs obtained after common de-replication between the two approaches. **(C)** Percentage of mapped reads on MAGs reconstructed by each approach, and on combined MAGs from both approaches, considering all mapping reads. MD = Metagenomic Distance, OR = Oceanic Region, MD+OR = MAGs from OR and MD approaches were combined together prior to reads mapping. **(D)** Prevalence-Abundance plot for MAGs reconstructed by both approaches (X-axis: MAG prevalence = number of metagenomic samples in which a MAG has a horizontal coverage above 0.3; Y-axis: MAG cumulative abundance. Percentage of mapped reads divided by the length of the MAG).

To evaluate the potential impact of co-assembly on assembly quality, we computed classical assembly quality metrics (N50 and L50) for both approaches. The metagenomic distance-based (MD) and the oceanic regions (OR) approaches actually reconstructed contigs of similar quality. No significant differences were detected in either the number of misassemblies, or the N50 and L50 metrics (Fig. S2). When considering the total number of bins generated following both co-assembly strategies, we found that both approaches reconstructed very similar numbers of bins: 10,748 bins generated using the MD approach, and 10,233 bins using the OR approach (Fig. 1B). To further compare both co-assembly strategies, as these bins may be very different in composition, we performed MAGs de-replication (37). The MD approach systematically reconstructed more MAGs than the OR approach, at both species (95% ANI) and strain (99% ANI) levels (supp. fig. S3B). Considering MAGs quality, medium quality (MQ) MAGs reconstructed by the MD approach were significantly more complete (Mann–Whitney U test, p=0.01, supp. fig. S4B), but evaluated as more contaminated (using checkM) than MQ MAGs reconstructed with the OR approach (Mann–Whitney U test, p=6.828 · 10^−05^, supp. fig. S4D). However, when considering the GUNC contamination metric (38), contamination levels observed in MQ MAGs of the MD approach were significantly lower than MQ MAGs of the OR approach (Mann–Whitney U test, p=2.352 · 10^−07^, supp. fig. S4F). Because GUNC assesses gene contamination based on all taxonomically annotated genes in a given genome, this latter approach may be considered as more robust than the checkM metric, and lead us to conclude that the MD approach actually reconstructed less contaminated MAGs. We found no significant differences in quality (completeness and contamination) for high quality (HQ) MAGs reconstructed by both approaches (supp. fig. S4). In addition, taxonomic annotations of strain-level de-replicated MAGs revealed a higher diversity recovered in the MD MAGs as compared to the OR MAGs (supp. fig. S5) in terms of number of distinct bacterial taxa, with a greater amount of annotated MAGs in MD (n=2006) as compared to OR (n=1869) MAGs.

Next, we also performed a global de-replication of MAGs in order to compare sets of MAGs recovered by both approaches at species and strain levels (see supp. Methods). Remarkably, we observed that both approaches reconstructed a very high number of exclusive MAGs (Fig. 1B). The OR approach reconstructed 575 species-level and 243 strain-level MAGs that were not recovered by the MD approach, while the latter did reconstruct 525 species-level and 323 strain-level MAGs that were not recovered by the OR approach. This result strongly emphasizes the influence of the co-assembly step prior to genome binning, in particular regarding how metagenomes are grouped for co-assembly. Given this observation, we aimed at determining which approach could captured a greater proportion of metagenomic diversity by back-mapping reads on MAGs generated by both approaches. While we observed a lower proportion of reads mapping to MD MAGs, as compared to OR MAGs, this proportion significantly increased when mapping on combined MAGs from both approaches. This result confirms that distinct and complementary MAGs are reconstructed using each approach. However, when only considering reads mapping to MAGs detected in samples, *i*.*e*. in which a given MAG has a minimum horizontal coverage (or breadth) of 30%, the MD approach recruited significantly more metagenomic reads as compared to the OR approach (Mann-Withney U test, p < 2.2 · 10^−16^, fig. 1D). Thus, although the OR MAGs were detected in more samples as compared to MD MAGs (Mann-Withney U test, p=4.6 · 10^−10^), the MD MAGs significantly improved the number and quality of reconstructed MAGs.

### Benchmarking assembly-binning strategies on simulated metagenomes

Different strategies for assembly and binning are currently used in the literature, each of them having its own advantages and disadvantages (14). Thus, we defined four assembly-binning strategies representing the most currently used approaches to reconstruct MAGs. Namely, we considered single-assembly (SA, i.e. the assembly of a single metagenome) and co-assembly (CA, i.e. the joint assembly of *n* metagenomes) approaches, as well as single-binning (SB, i.e. genome binning solely using (co-)abundance information from metagenome(s) used to perform the (co-) assembly) and co-binning (CB, i.e. genome binning using co-abundance information from all metagenomes) approaches. We thus evaluated the following four strategies: Single-Assembly with Single-Binning (SASB), Single-Assembly with Co-Binning (SACB), Co-Assembly with Single-Binning (CASB), and Co-Assembly with Co-Binning (CACB). We compared the performances of these four strategies on three different datasets, the CAMI (15) high complexity dataset, a lower complexity mockdataset generated using CAMISIM (39), and a human microbiome dataset from the Human Microbiome Project (40).

First, to evaluate and compare these four strategies on simulated metagenomes, we applied our MD clustering algorithm on the CAMI high-complexity dataset (15). The CAMI high-complexity dataset is composed of five metagenomic samples simulated from a community of 596 known reference genomes and 478 circular elements. The optimal solution identified for the co-assembly regrouped all five metagenomes, probably due to the small number of metagenomes (n=5) and the fact that they were simulated from the same pool of reference genomes. Therefore, only one co-assembly (of all 5 samples) was performed, and the CACB and CASB strategies were thus equivalent. Following genome binning using MetaBAT2 (9), the SACB strategy reconstructed the highest number of bins (> 400 genome bins), while the CASB and SASB strategies reconstructed about 300 and 200 bins, respectively (Fig. 2A).

**FIG 2.**
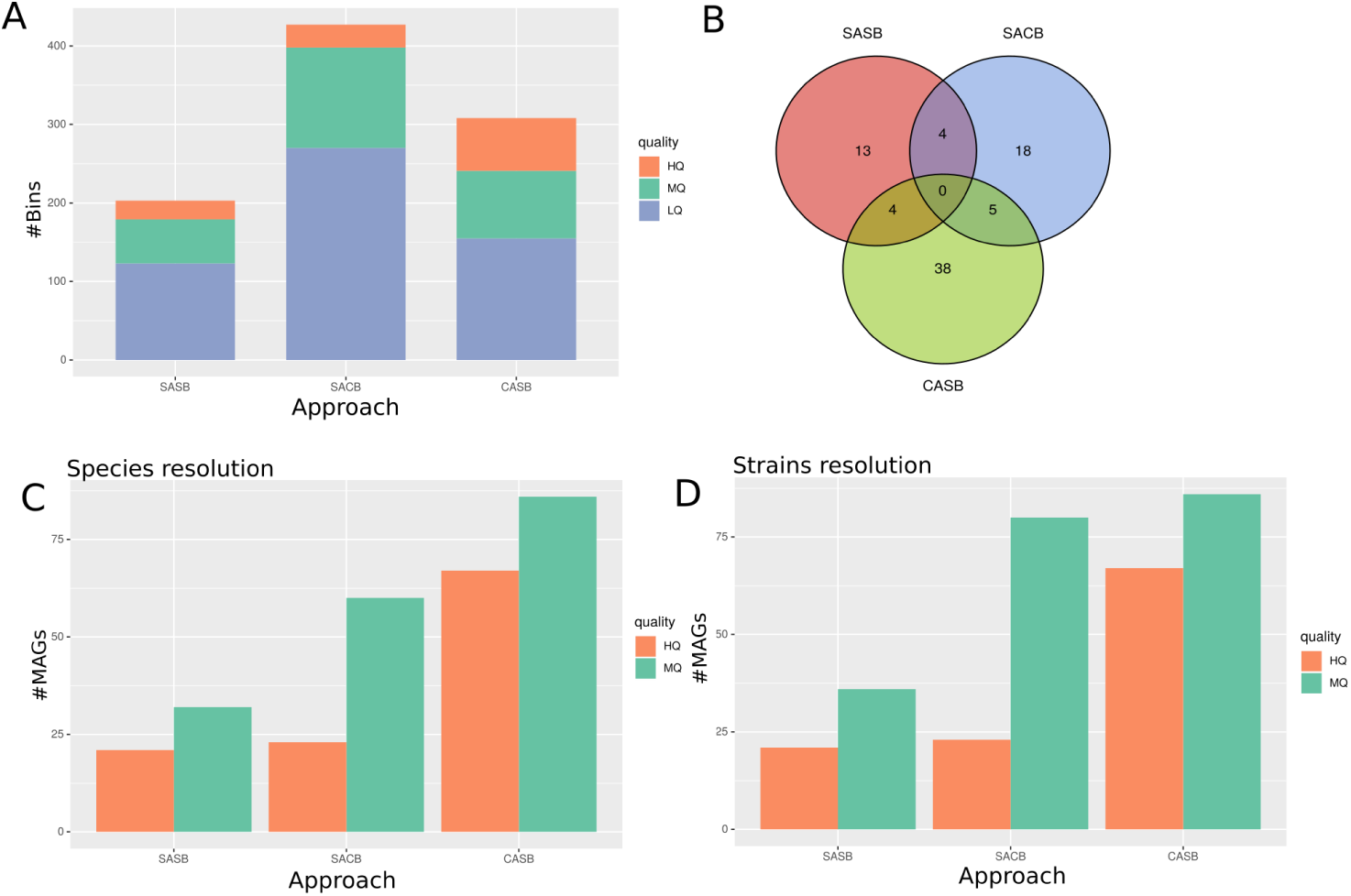
Evaluating assembly-binning strategies on the CAMI dataset. **(A)** Total number of bins obtained after binning step. Colours represent quality of genome bins estimated using CheckM: High Quality (HQ), Medium Quality (MQ), and Low Quality (LQ). **(B)** Number of MAGs mapping to a source genome within each strategy, corresponding to the number of expecting genomes in the set of MAGs of each strategy. The diagram represents thus the common genomes found in each strategy. **(C**,**D)** Number of reconstructed MAGs after independent de-replication using dRep for each binning strategy, at **C** Species resolution, consisting in a 95% ANI score de-replication; and **D** Strains resolution, consisting in a 99% ANI score de-replication. SASB: Single-Assembly-Single-Binning, SACB: Single-AssemblyCo-Binning, CASB: Co-Assembly-Single-Binning.

After de-replication, we compared the MAGs obtained for each strategy to the CAMI reference source genomes. When considering the distribution of expected genomes across all three strategies, we observed that the CASB strategy reconstructed more expected genomes than both single-assembly strategies (SASB and SACB). Surprisingly, we did not find expected genomes common to all strategies (Fig. 2B), which highlights the actual complementarity of these strategies. When considering only de-replicated genomes, CASB produced the highest number of MAGs. This difference was clear for HQ MAGs, for which CASB produced about 2.5 times more MAGs as compared to single-assembled strategies, with both SACB and SASB generating a comparable number of HQ MAGs (Fig. 2CD).

The number of reconstructed MAGs was also dependent of the de-replication level. At strain level, both single-assembly approaches reconstructed more non-redundant MAGs compared to species level, while CASB reconstructed the same number of MAGs at both species and strain levels. However, this increase concerned only the MQ MAGs, as the number of HQ MAGs remained unchanged (Fig. 2D). We did not find any significant differences in MAGs completeness between the different strategies, considering either HQ MAGs or MQ MAGs (supp. fig. S6 A&B). However, we did observe differences in contamination estimated from Single-Copy Genes (SCGs) using checkM. CASB HQ MAGs were less contaminated than SASB (Mann-Withney U test, p=0.01) and SACB (Mann-Withney U test, p=0.04) HQ MAGs, while SACB MQ MAGs were less contaminated than SASB MQ MAGs (Mann-Withney U test, p=0.03) (supp. fig. S6 C&D). When considering MAGs contamination estimated using taxonomically annotated genes using GUNC, CASB MAGs were predicted most contaminated (supp. fig. S6 E&F), with significant differences observed with either SASB (Mann-Withney U test, p=3 · 10^−4^) and SACB (Mann-Withney U test, p=2 · 10^−4^) MAGs. We did not find any other differences in contamination levels between the four strategies.

Given that the MD clustering approach did not identify optimal clusters to co-assemble within the CAMI dataset, we used CAMISIM (39) to simulate an additional metagenomic dataset with a higher number of samples and a lower complexity. We thus simulated 20 metagenomes with a similar diversity of 100 reference genomes. On this simulated dataset, the MD clustering approach identified 8 optimal clusters to co-assemble. Here, the co-assembly-based strategies (CACB and CASB) reconstructed more bins than the single-assembly-based strategies (3A), also when considering only HQ bins. After de-replication, we aimed to identify expected genomes among recovered MAGs by mapping them to reference genomes used for the metagenomes simulation. The majority (n=29) of expected genomes we identified were reconstructed in all four strategies (3B). The SACB strategy recovered a short majority of expected genomes (n=33), as compared to CACB ans SASB (n=32), and CASB (n=31). However, the number of de-replicated MAGs was higher for both co-assembly strategies compared to single-assembly strategies (3C).

The drop in de-replicated MAGs from single-assembly strategies is likely a consequence of the higher number of assemblies performed in both SASB and SACB strategies. As single-assemblies are more numerous than co-assemblies, there is thus a higher probability to reconstruct, independently, several times the same MAG. Finally, using this simulated dataset, we did not detect any significant differences in the quality of MAGs reconstructed by the four strategies, neither in their completeness nor in their contamination levels (supp. fig. S7).

### Comparing assembly-binning strategies on real metagenomes

To further compare the four genome reconstruction strategies, we applied them to a real metagenomic dataset, which is more complex in terms of species diversity and composition. Human gut microbiome studies represent a large fraction of publicly available metagenomes and are also good case studies as they represent metagenomes with intermediate complexity compared to soil or ocean metagenomes. Thus, we focused on analysing a selection of 150 metagenomes of human gut microbiomes from the Integrative Human Microbiome Project (HMP) (40).

Here, the MD-based clustering approach identified 64 metagenomic clusters to co-assemble. When comparing all four strategies before de-replication, both single-assembly strategies reconstructed more genome bins than both co-assembly strategies (4A). Next, in order to determine how many MAGs we could expect to reconstruct at best by each strategy, we de-replicated altogether genome bins reconstructed by all strategies. The resulting number of de-replicated MAGs thus represents the highest number of MAGs we would be able to reconstruct with the HMP dataset combining all four strategies. We then compared each strategy by considering what proportion of the maximum number of MAGs each strategy was able to reconstruct (4B). After de-replication at the species level, despite the fact that single-assembly strategies recovered more bins, we observed that both co-assembly strategies reconstructed more MAGs than single-assembly strategies. Also, for both co-assembly and single-assembly strategies, the co-binning actually allowed to reconstruct more MAGs than the single-binning approach (4B&C), which underlines the importance of integrating cross-samples information when binning genomes. However, after de-replication at strain level, we observed that the SACB strategy reconstructed more MAGs than CASB, while the SASB strategy reconstructed more HQ MAGs than the CASB strategy (4B&D).

We also compared the MAGs quality (completeness and contamination) produced by each assembly-binning strategy. Differences in completeness were only observed between the SACB and CASB strategies, with SACB HQ MAGs being more complete than CASB HQ MAGs (supp. fig. S8A). Here, we also used both checkM (SCG-based) and GUNC (taxonomy-based) complementary approaches to estimate contamination. GUNC was able to detect more subtle differences in contamination between strategies than the checkM algorithm (supp. fig. S8). These observed differences demonstrated that co-binning strategies actually produced less contaminated MAGs than single-binning strategies, at all MAGs quality levels. Overall, these distinct results when de-replicating MAGs at species or strain level suggest that no single strategy can fit all needs. Therefore, the choice of an assembly-binning strategy should be informed by a biological question and considering the microbiome complexity under study.

### Comparing MAGNETO to similar metagenomics worklows

Finally, we compared the performances of MAGNETO to metagenomics workflows dedicated to MAGs reconstruction, namely METAWRAP (22), ATLAS (24) and nf-core/mag (25). We chose these three tools as they use similar softwares to perform assembly and binning, namely MEGAHIT (41) and MetaBAT2 (9). The comparison of the workflows was performed using the HMP dataset. ATLAS is a workflow only permitting single-assembly of metagenomes, but integrates a binning refinement module using DAStool (42), which constitutes a good opportunity to evaluate whether single-assembly could perform better after binning refinement. METAWRAP also contains a binning refinement module, albeit less complex than the DAStool methodology. This refinement module performs pairwise alignment of MAGs to detect redundant genomes, to then only conserve MAGs showing the best quality amongst detected duplicated MAGs. nf-core/mag uses the exact same tools as our workflow to perform assembly and binning. As compared to ATLAS, we observed that MAGNETO systematically reconstructed more MAGs using any of the four assembly-binning strategies (Table 1). However, it also reconstructed less MAGs than METAWRAP. The higher number of MAGs produced by METAWRAP may be explained by its refinement module coupling several binners, as these binners may reconstruct more non-redundant MAGs, thus increasing their numbers. However, MAGNETO and nf/core-mag reconstructed the same number of MAGs for both CASB or CACB strategies. These similar results are most likely explained by the absence of a bins refinement module, and by the fact that in both workflows, the binning step used the exact same parameters.

**TABLE 1.** Number of reconstructed MAGs for the HMP dataset. Comparison of the number of MAGs reconstructed with different workflows, and different strategies, after dereplication at strains resolution. MAGs: number of dereplicated MAGs; HQ: High Quality (Completeness > 90%, Contamination < 5%), MQ: Medium Quality (Completeness > 50%, Contamination < 10%).

| Pipeline | Strategy | MAGs |  |
| --- | --- | --- | --- |
|  |  | HQ | MQ |
| ATLAS |  |  |  |
|  | SASB | 253 | 120 |
| METAWRAP |  |  |  |
|  | SASB | 302 | 295 |
|  | SACB | 320 | 242 |
|  | CASB | 377 | 320 |
|  | CACB | 386 | 350 |
| nf-core/mag |  |  |  |
|  | CASB | 277 | 261 |
|  | CACB | 361 | 300 |
| MAGNETO |  |  |  |
|  | SASB | 280 | 251 |
|  | SACB | 311 | 286 |
|  | CASB | 277 | 261 |
|  | CACB | 361 | 300 |

### Design and implementation

MAGNETO is a Snakemake (43) workflow connecting open-source bioinformatics software, all available from Bioconda and conda-forge. Snakemake was chosen for its flexibility, its capacity to run both locally and on clusters, and its Conda management automating software installation. MAGNETO includes several tools designed for metagenomic studies. First, reads trimming is performed using fastp (44) and FastQ Screen (45). The co-assembly module relies on Simka (35), which estimates metagenomic distances between samples based on their k-mers composition. MEGAHIT (41) then performs reads assembly/co-assembly. We use MetaBAT2 (9) to bin contigs, and we assess the quality of bins using CheckM (46). The de-replication of bins into MAGs (bins of at least high of medium quality) is performed using dRep (37). Notably, MAGNETO can also be used to establish gene catalogs, to better capture metagenomic gene diversity by producing a non-redundant set of genes through sequence clustering at a user-defined sequence identity cutoff (e.g. 95%) using Linclust (47). GTDB-tk (48) is used to perform taxonomic annotation of dereplicated MAGs, and eggNOG-mapper (49) is used to perform the functional annotation of MAGs as well as the gene catalog (see Figure 5).

**FIG 3.**
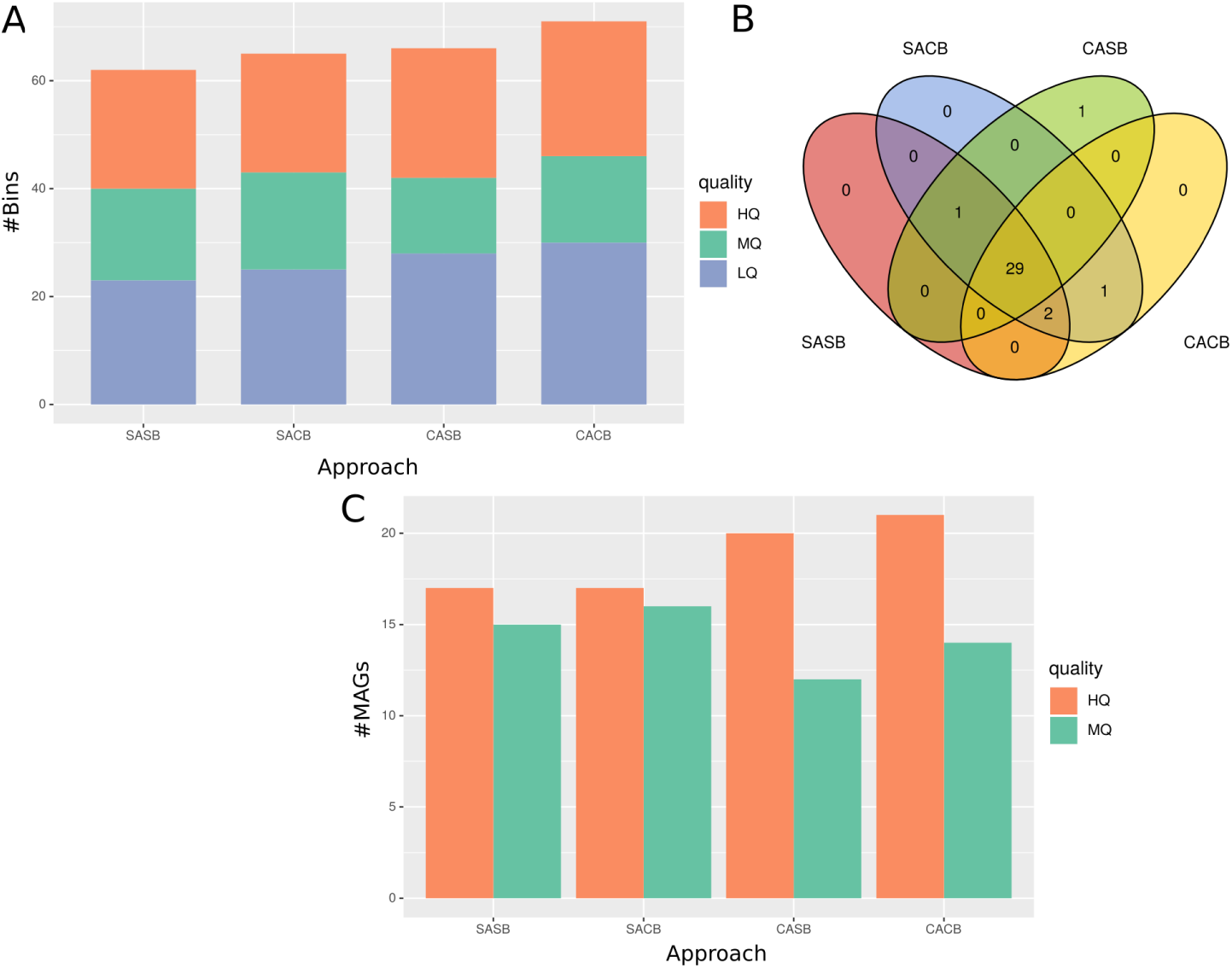
Evaluating assembly-binning strategies on simulated metagenomes. **(A)** Total number of bins obtained after the binning step. Colours represent quality of genome bins estimated using CheckM: High Quality (HQ), Medium Quality (MQ), and Low Quality (LQ). **(B)** Number of source genomes found in each strategy. Each number represents the number of times a MAG from a strategy maps against a source genome. Intersections represent common genomes between strategies. **(C)** Number of de-replicated MAGs obtained, after independent de-replication by dRep for each strategy. As the genomes are all represented with one single strain, de-replication at both species or strain resolution gives the same number of de-replicated genomes, so only one de-replication resolution is shown.

**FIG 4.**
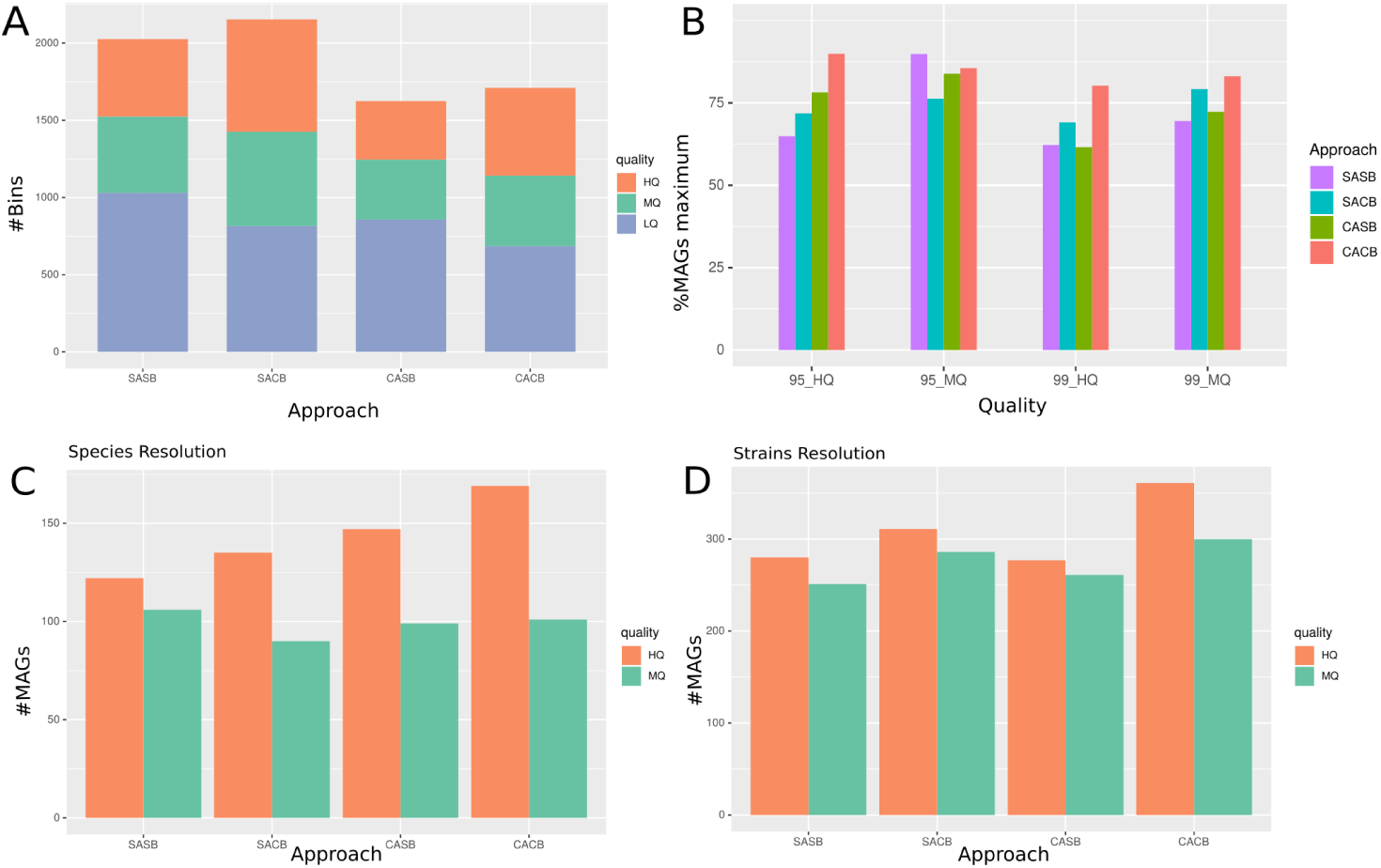
Evaluating the binning strategies on the HMP dataset. **(A)** Total number of bins reconstructed per strategy. Colours represent the MAGs qualities, estimated with CheckM. **(B)** Proportion of MAGs reconstructed for each strategy, after common de-replication of the four strategies, at the species resolution (95% identity) or at the strain resolution (99% identity). Number of de-replicated MAGs from each strategy is compared to the number of maximum expected MAGs, which is the number of MAGs obtained after dereplication of all the four strategies together. **(C**,**D)**: Number of reconstructed MAGs after independent de-replication using dRep for each binning strategy, at **C** Species resolution, consisting in a 95% ANI score de-replication; and **D** Strains resolution, consisting in a 99% ANI score dereplication. SASB: Single-Assembly-Single-Binning, SACB: Single-Assembly-Co-Binning, CASB: Co-Assembly-Single-Binning, CACB: Co-Assembly-Co-Binning. HQ: High Quality, MQ: Medium Quality, LQ: Low Quality.

**FIG 5.**
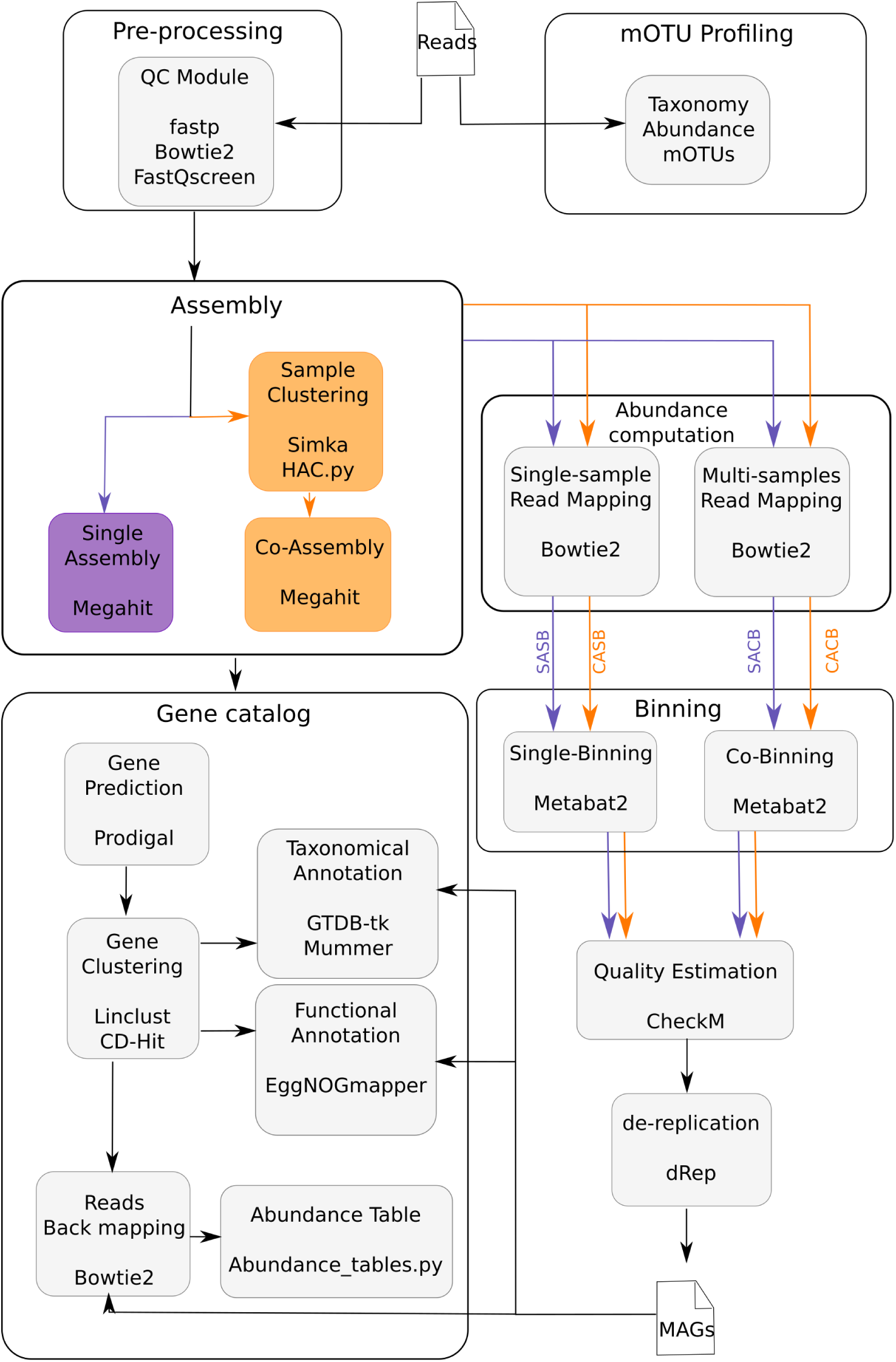
Overview of the MAGNETO workflow. Summary view of modules implemented in the MAGNETO workflow, with the name of the software or script associated to each task. The workflow can be launched for a complete run, to process raw reads into a gene catalog and MAGs, but each module can also be run independently. In purple: path to perform single-assembly, corresponding to SASB (Single-Assembly-Single-Binning) and SACB (Single-Assembly-Co-Binning) strategies, and orange: path to perform co-assembly, corresponding to CASB (Co-Assembly-Single-Binning) and CACB (CoAssembly-Co-Binning) strategies.

A more complete description of each module implemented in MAGNETO is available in the Material and Methods section. The four binning strategies are directly configurable by the user, and a quick configuration allows to perform from a single to all strategies for reconstructing MAGs. Notably, MAGNETO is currently the unique workflow providing an automated approach to define clusters of metagenomes for co-assembly. Importantly, MAGNETO and nf-core/mag are also the only workflows allowing users to perform a co-binning strategy. A synthetic comparison of functionalities provided by the workflows tested in this study are available in Table 2.

**TABLE 2.** Comparison of tasks performed by evaluated workflows.

| Steps | ATLAS | METAWRAP | nf-core/mag | MAGNETO |
| --- | --- | --- | --- | --- |
| **Pre-processing** |  |  |  |  |
| Reads Trimming | ✓ | ✓ | ✓ | ✓ |
| Contamination | ✓ | ✓ | ✓ | ✓ |
| **Assembly** |  |  |  |  |
| Co-assembly possible |  | ✓ | ✓ | ✓ |
| Compute sets to co-assemble |  |  | ✓ |  |
| **Binning** |  |  |  |  |
| Co-binning possible |  | ✓ | ✓ | ✓ |
| Multiple binning software | ✓ | ✓ |  |  |
| Bin refinement | ✓ | ✓ |  |  |
| Bin reassembly | ✓ | ✓ |  |  |
| **Post-processing** |  |  |  |  |
| MAGs quality check | ✓ | ✓ | ✓ | ✓ |
| de-replication step | ✓ | ✓ | ✓ | ✓ |
| Genome annotation | ✓ | ✓ | ✓ | ✓ |
| Gene catalogue |  |  | ✓ | ✓ |
| **Reproducibility** |  |  |  |  |
| Workflow Management | ✓ |  | ✓ | ✓ |
| Packages management | ✓ |  | ✓ | ✓ |

## DISCUSSION

In this work, we present MAGNETO, a fully automated workflow enabling genome-resolved metagenomics. It implements a novel approach to compute clusters of metagenomes for co-assembly without *a priori* knowledge, as well as complementary assembly-binning strategies to maximize MAGs recovery towards specific goals. MAGNETO also provides key functionalities, from the construction and annotation of gene catalogs, to the generation of genes and genomes abundance matrices.

### An unsupervised approach to metagenomic co-assembly

We demonstrated the utility of a non-supervised metagenomic-distance based approach to guide metagenomics co-assembly on a large set of ocean metagenomes. Indeed, clusters of metagenomes identified by the MD-based approach did not overlap with oceanic regions previously used for guiding co-assembly of these metagenomes (13). As anticipated, this implies that, in the ocean, geographic distances do not necessarily reflect compositional metagenomic distances between microbial communities. This observation can likely be explained by the fact that the composition of marine microbial communities are significantly structured through environmental filtering by key abiotic factors such as temperature (28) and ocean currents influencing species dispersal (50). Interestingly, the MD-based clustering analysis grouped together in a single cluster (cluster #1, see Figure 1A) metagenomes from sampling stations facing upwelling currents. As upwelling regions are influenced by deep ocean currents raising cold nutrient-rich waters to the surface, they can significantly impact species diversity of marine microbial communities towards richer states (51, 52).

The rationale behind our metagenomic distance-based approach to perform coassembly was to infer which metagenomes should be grouped together in an unsupervised fashion without a priori knowledge. The aim was to develop an approach that could guarantee the actual closeness of the metagenomes to co-assemble, thus emphasizing the increase in species-specific reads abundance for the assembler. Although the co-assembly of closely related metagenomes have been shown to erode contigs quality (15, 12), we could show that our approach did not increase fragmentation or misassemblies within contigs (supp. fig. S4). In fact, our MD approach reconstructs MAGs that are more complete, and less contaminated than the OR approach (supp. fig. S4). Although both metrics we used to estimate MAGs contamination reported contradictory results, we argue that GUNC (38) likely provides better estimates of contamination as it is based on a much larger set of genes as compared to CheckM (46), which assess contamination solely based on SCGs. As SCGs represent highly-conserved genes across all taxa, co-assembling similar metagenomes may actually increase the probability to assemble or bin core regions of closely related genomes. A higher fragmentation of the genomes was already observed following the co-assembly of metagenomes with closely related strains (15, 53), although it was also shown not to affect completeness nor the contamination of co-assembled genomes (37). Accessory regions may thus be less affected by co-assembly, although they are also generally more difficult to bin (12).

We observed a very high number of exclusive MAGs between the OR and MD approaches, namely 525 for MD and 575 for OR, representing 31.2% and 33.3% of the MAGs reconstructed by each approach, respectively (Fig. 1B). This result indicates that, even if our approach performs better in terms of reconstructed MAGs quality, it nevertheless does not capture the same information from metagenomes as compared to the OR approach. This is confirmed by the increase in proportion of recruited reads when back-mapping to combined MAGs from both approaches (Fig.1C). Thus, combining the MD approach with a co-assembly based on *a priori* knowledge (when available) may represent a good opportunity to better capture the actual bacterial diversity in metagenomes. However, the proportion of mapped reads was significantly higher on MD MAGs as compared to OR MAGs when considering only detected MAGs in samples (Fig.1D). Here, we could show that the OR approach reconstructed MAGs recruiting a higher proportion of reads, but that this higher proportion was mainly driven by MAGs displaying a very low horizontal coverage (< 30%), suggesting these MAGs contained relatively small genomic regions recruiting a high proportion of reads. These observations, coupled with the smaller contamination observed in OR MAGs when estimated using SCGs, may imply that the OR approach allows a better reconstruction of core genomic regions, which are shared among a higher proportion of organisms.

Applying the MD-based co-assembly approach on the HMP dataset, we found that the identified clusters of metagenomes mostly corresponded to the IBD pathology affecting the patients (supp. fig. S9). Indeed, a majority of clusters containing metagenomes from healthy patients did not contain any metagenomes related to IBD (16 out of 23 clusters contain non-IBD metagenomes), and a majority of the clusters containing CD or UC patients are composed of metagenomes associated with only the same type of IBD (26 out of 34 clusters contain IBD metagenomes). This observation emphasizes the relevance of our method, as changes in the composition of the gut microbiota have been associated with IBD diagnostic (40, 54, 55).

### A systematic comparison of assembly-binning strategies

When comparing the four different assembly-binning strategies we defined herein, we observed that *co*strategies systematically reconstructed more MAGs than *single*-strategies. Notably, the CACB strategy was identified as the best performing in terms of number of recovered MAGs, across all (simulated and real) datasets we considered. This may be explained by i) the increase in (rare) reads abundance through the co-assembly, and ii) the higher amount of co-abundance information integrated into the co-binning process (7, 26). On simulated datasets, co-assembly strategies systematically reconstructed more MAGs after de-replication, while applying single-binning or co-binning. However, this was not the case when analysing the HMP dataset, for which the SACB strategy reconstructed more strain-level MAGs than CASB. This may be due to an uneven distribution of strains across metagenomes. Indeed, human gut microbiomes tend to be personal and usually exhibit higher inter- than intra-individual community variations at strain level (56, 57). Overall, if gut strains are individual-specific and thus only occur in a low number of metagenomes, co-assembly will be less effective to actually increase strain-specific reads for improving their assembly. This result actually suggests that an MD-based approach integrating single-nucleotide polymorphism (SNP) information would be useful to improve the reconstruction of strain-level MAGs.

### A multi-sample assembly-binning strategy maximizes genomes recovery

We showed that co-assembly approaches usually reconstructed higher numbers of (MQ) MAGs, albeit with a tendency to be more contaminated (HQ MAGs). As previously reported (12), this underlines the utility of co-assembly to recover rare or less-abundant genomes, and to maximise MAGs recovery from a limited number of metagenomes. Here, co-binning strategies (SACB & CACB) systematically reconstructed less contaminated MAGs than single-binning strategies (SASB & CASB) in datasets for which differences in MAGs quality could be detected between strategies. Thus, multi-sample co-abundance information computed across a minimum number of metagenomes appears particularly relevant to improve genome binning and to limit the erroneous grouping of contigs. However, the co-binning strategy may represent a severe limitation as it requires larger computational resources (CPU time and disk space), since it implies performing *N* ^2^ reads mapping operations, with *N* the total number of metagenomes. For the CAMI dataset, differences in MQ MAGs quality between strategies were in contradiction with analyses of the other datasets, although the HQ MAGs comparison pointed towards similar conclusions as in the other datasets. This may be explained by the different number of MAGs reconstructed between each strategy. The 80 MAGs reconstructed by the SASB strategy may belong to abundant organisms, thus implying a smaller risk to increase contamination. However, as SACB and CASB reconstructed almost twice the number of MAGs compared to SASB, the MQ MAGs recovered by these strategies may belong to less abundant genomes, hence these MAGs may be harder to reconstruct with a few samples (n=5), and thus may be more prone to contamination.

Interestingly, the effect of the co-assembly step on MAGs contamination is unclear. So far, only a few methods, including CheckM and GUNC, exist to estimate MAGs quality. When considering CheckM on the HMP dataset, single-assembly strategies reconstructed less contaminated MAGs than co-assembly strategies. However, when considering contamination estimated by GUNC, co-assembly strategies constructed less contaminated MAGs. These results underline the crucial need to develop more accurate methods to properly estimate MAGs quality, and also highlight the utility to confront methods using complementary strategies to estimate genome quality.

Co-assembly constitutes a useful and affordable strategy for shallow sequenced metagenomes or when the number of metagenomes to co-assemble is limited. In such cases, the increase in complexity of the assembly is limited, thus removing the main computational limitation of co-assembly. Similar to co-assembly, co-binning is also impacted by metagenomic sequencing depth, as the computation time obviously increases with the number of reads. As demonstrated, the co-binning strategy represents a powerful and useful, though computer-intensive, strategy when numerous samples are available, as it helps to reconstruct more HQ MAGs. A potential perspective for improving the co-binning process would be to identify an optimal number of samples to compute co-abundances in order to optimize its cost-benefit ratio.

## MATERIALS AND METHODS

### Reads pre-processing

Raw reads were filtered using fastp (44) and FastQ Screen (45). fastp filters reads on their quality, length and complexity. FastQ Screen is a tool allowing to control contamination within metagenomic samples, by mapping their reads to reference genomes. These two tools provide results reports to the user that are useful to evaluate reads quality.

### Assembly

We performed reads assembly using MEGAHIT (41), as this assembler provides an excellent trade-off between computational requirements and assembly quality (58). metaSPAdes (59) could have been considered as it provides better performances than MEGAHIT in terms of overall percentage of the metagenome recruited in the assembly (60) and maximum length of scaffolds produced (60, 61). However, this performance increase occurs at the cost of a greater consumption of computational resources (58) and a presence of a greater proportion of misassembled sequences in contigs than MEGAHIT (60). More importantly, metaSPAdes was originally not designed to perform co-assembly, which constituted a major drawback in our workflow. Moreover, MEGAHIT is able to capture micro-diversity from the metagenomes more efficiently than metaSPAdes, as it discards less low-abundant reads during assembly (58). Co-assemblies of the marine metagenomes were performed using the *–presets metalarge* option, as these metagenomes revealed to be highly complex. All other assemblies were performed using the *–preset meta-sensitive* option.

### Co-assembly strategy

In order to determine which samples to co-assemble, we used Simka, a de novo and scalable tool for comparative metagenomics (35). Simka computes different distances based on *k* -mer counts, instead of species counts. In our case, we used their modified Jaccard (or *AB* -Jaccard) distance rather than the default Bray-Curtis distance, as the latter does not satisfy triangle inequality. Once the distance matrix from Simka was computed, samples were then clustered using a Ward-based hierarchical agglomerative clustering (62). Then, we iteratively cut the dendrogram and assessed partitioning quality using the Silhouette method (36).

### Genome binning strategies

Binning was performed using MetaBAT2 (9), as it is currently one of the fastest and best performing genome binner. We set the minimum length for contigs to be binned to 1500 nucleotides. As MetaBAT2 uses composition and abundance to perform binning, a preliminary step to map reads back to assembled contigs was performed to measure abundance. Reads mapping was achieved using Bowtie2 (63).

Instead of computing an abundance metric only from the metagenome assembled into contigs, MetaBAT2 may compute a co-abundance metric using contig coverage from several samples, even if these samples do not participate in the assembly. A co-abundance metric computed from several samples increases the quality of the genome bins produced (64). Depending on the number of samples used to compute contigs abundance, the corresponding metric is either an abundance or a co-abundance metric. Thus, two strategies can be pursued in order to perform binning: (i) single-binning, which uses abundance of contigs measured from assembled metagenome(s); or (ii) co-binning, which uses co-abundance of contigs measured from all the metagenomes of a dataset. Combined with the decision of performing either single-assembly or co-assembly, we defined four binning strategies: Single Assembly of one metagenome with Single-Binning (SASB), Single Assembly of one metagenome with Co-Binning (SACB), Co-Assembly of one set of metagenomes with Single-Binning (CASB), and Co-Assembly of one set of metagenomes with Co-Binning (CACB).

### Genome bins quality

Genome bins quality was defined by two metrics, namely completeness and contamination. Completeness measures the fraction of the initial genome captured, while completeness measures the fraction of alien sequences; both rely on the presence-absence patterns of universal Single-Copy marker Genes (SCGs). To assess genome bins quality, we used CheckM (46) and GUNC (38). Based on contamination and completeness, we distinguished three standard quality levels for bins (65) : (i) high-quality bins (HQ) with completeness > 90% and contamination < 5%, (ii) medium-quality bins with completeness > 50% and contamination < 10%, while the (iii) low-quality bins (LQ) are bins that are neither HQ nor MQ. Only HQ and MQ bins were then considered to be MAGs. The comparisons of reconstructed MAGs quality from different strategies were performed using Mann-Withney U test using R (66). As a MAG may be reconstructed independently either in two (or more) samples or two (or more) co-samples, MAGs are also de-replicated using dRep (37). Two MAGs were considered to be duplicated if their pairwise ANI (Average Nucleotide Identity) score was above a given identity threshold *t*, (*t* being a percentage of sequence identity) on more than 60% of their bases (67). We consider two different values for *t* : *t* = 0.95, which corresponds to a de-replication at species level, and *t* = 0.99, which corresponds to a de-replication at strain level (68).

### Genome annotation module

Functional and taxonomic annotations were performed for the strain-level MAGs collection, which encompasses the species-level collection. To perform functional annotation of MAGs, we used eggNOG-mapper (49), and we used GTDB-tk (48) to perform taxonomic annotation. Finally, reads of each sample are mapped back onto both species- and strain-level MAGs collections using Bowtie2, and an abundance table is produced using an in-house Python script.

### Gene annotation module

Coding DNA Sequences (CDS) are detected on assembled contigs per-sample (single-assembly) using Prodigal (69). Genes from all samples are clustered at 95% identity using Linclust (47) in order to produce a non-redundant set of genes (gene95 collection). EggNOG (49) and MMSEQ2 (70) are used to annotate this gene collection, for functional and taxonomic information, respectively. Finally, reads from each sample are mapped back onto the gene95 collection using Bowtie2, and an abundance table is produced.

### Datasets

The marine metagenomes dataset corresponds to the same 93 oceanic metagenomes as processed in Delmont et al. (13), which are available at the European Bioinformatics Institute (EBI) repository under project ID ERP001736. In order to benchmark the assembly-binning strategies, we simulated two different mock metagenomes datasets. First, we used CAMI (15) high-complexity dataset, which is a 75-Gbp time series dataset sequenced into short reads, composed of five samples from a high-complexity community with correlated log normal abundance distributions (596 genomes and 478 circular elements). However, the above dataset did not allow us to assess our clustering method, which was not able to determine an optimal clustering to perform CASB and CACB. Thus, a simple co-assembly strategy, gathering all 5 samples, was performed on this dataset and was referred as a CASB strategy. We also simulated two more datasets, composed of more samples than CAMI dataset, in order to better evaluate our co-assembly module. The first dataset we simulated was composed of 10 metagenomes containing different abundances of 20 genomes, with each metagenome sizing 1 Gbp. Our second dataset was composed of 20 metagenomes containing different abundances of 100 genomes, each sample sizing 1 Gbp. Both these simulated datasets were generated using CAMISIM (39) with each metagenome’s size fixed to 1 Gbp. Each metagenome contained the 100 source genomes, with specific abundance distribution created by sampling from a log-normal distribution with *μ* set to 1 and *σ* to 2 (default values). To simulate the first mock dataset, we used the 20 test genomes used by default in CAMISIM. The 100 genomes used for the simulation of the second mock dataset were randomly sampled from CAMI high-complexity source genomes. Other CAMISIM parameters were set to their default values.

For the human gut microbiome dataset, we collected raw data from the Integrative Human Microbiome Project (iHMP) (40). Based on the availability of both metagenomics and metatranscriptomics data, we selected a subset of 150 metagenomes, representing 80 individuals and patients followed during one year. Metagenomes were extracted from stool samples and were sequenced using Illumina technology. The samples subset we analysed here was composed of three sample groups of strictly equal size characterized by diagnosis, with individuals with no Inflammatory Bowel Disease (non-IBD), and patients either diagnosed with Crohn’s disease (CD) or Ulcerative Colitis (UC).

## ACKNOWLEDGMENTS

We wish to thank the CNRS MITI through the interdisciplinary program Modélisation du Vivant (GOBITMAP grant to SC), and the H2020 European Commission project AtlantECO (award number 862923). The funders had no role in study design, data collection and interpretation, or the decision to submit the work for publication.

Computational support was provided by the bioinformatics core facility of Nantes (BiRD, Biogenouest), University of Nantes, France.

B.C. and S.C. designed the study. B.C. and M.M. performed the experiments. B.C., M.M., A.B., G.F. and S.C. analyzed the data and interpreted the results. B.C., G.F. and

S.C. wrote the paper, with input from M.M. and A.B.. We declare that we have no competing interests.

## SUPPLEMENTARY MATERIAL

### Supplementary Methods

The construction of the Fig. 1B panel implied a global de-replication of the *TARA* Oceans MAGs, performed by combining sets of MAGs recon-structed with both approaches using dRep as described in the methods section. Once this global de-replication was performed, the total number of de-replicated MAGs (dMAGs) related to one approach is thus:

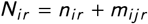

with *n* the number of dMAGs already reconstructed by the approach *i*_*i*_, and *m*_*ij*_ the number of dMAGs reconstructed by approach *j*, but located in a de-replication cluster containing at least 1 MAG reconstructed by the approach *i*, at the de-replication resolution *r*. From the dRep output, we can identify the de-replication cluster each MAG belongs to, and the number of members located in the same de-replication cluster. We can then list, for a given de-replication resolution *r*, the set of dMAGs related to one approach *i*, searching for each non-unique dMAG (*i*.*e*. having at least 1 neighbour in its de-replication cluster) of the approach *j*, if there is at least one MAG from the approach *i*. Thus, to detect shared dMAGs between both approaches, we identified the common elements between the four sets of *N*_*i r*_ dMAGs.

To detect shared genomes between sets of genomes related to different de-replication resolutions, we should precise that the set of dMAGs at species-level for one approach is completely included in the set of dMAGs at strain-level from this same approach. Thus, we can non-ambiguously identify clustering relationships between two MAGs from different de-replication levels, or find exclusive MAGs reconstructed by one approach, but present at both species and strain levels.

**FIG S1.**
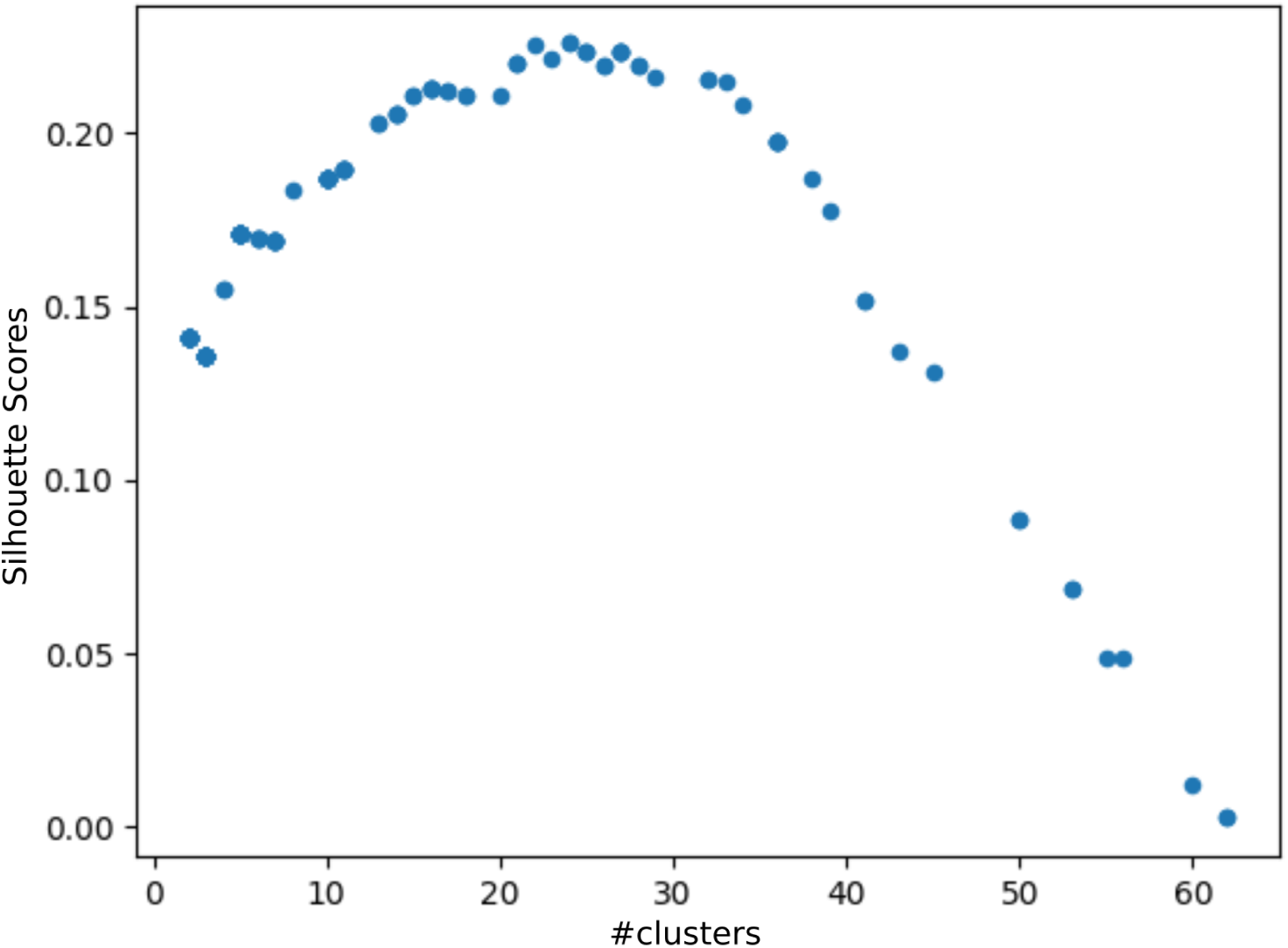
Identifying metagenomic distance-based optimal clusters for *TARA* Oceans metagenomes. Silhouette scores obtained when clustering the metagenomes into *k* groups. X-axis: values of *k* tested, *k* varying from 2 to *n* − 1, with *n* = 93 (the total number of samples used). Y-axis: Values of the Silhouette score estimated for value *k* tested. A high Silhouette score yields a better clustering. A maximum score is observed for *k* = 24, which thus represents the optimal clustering.

**FIG S2.**
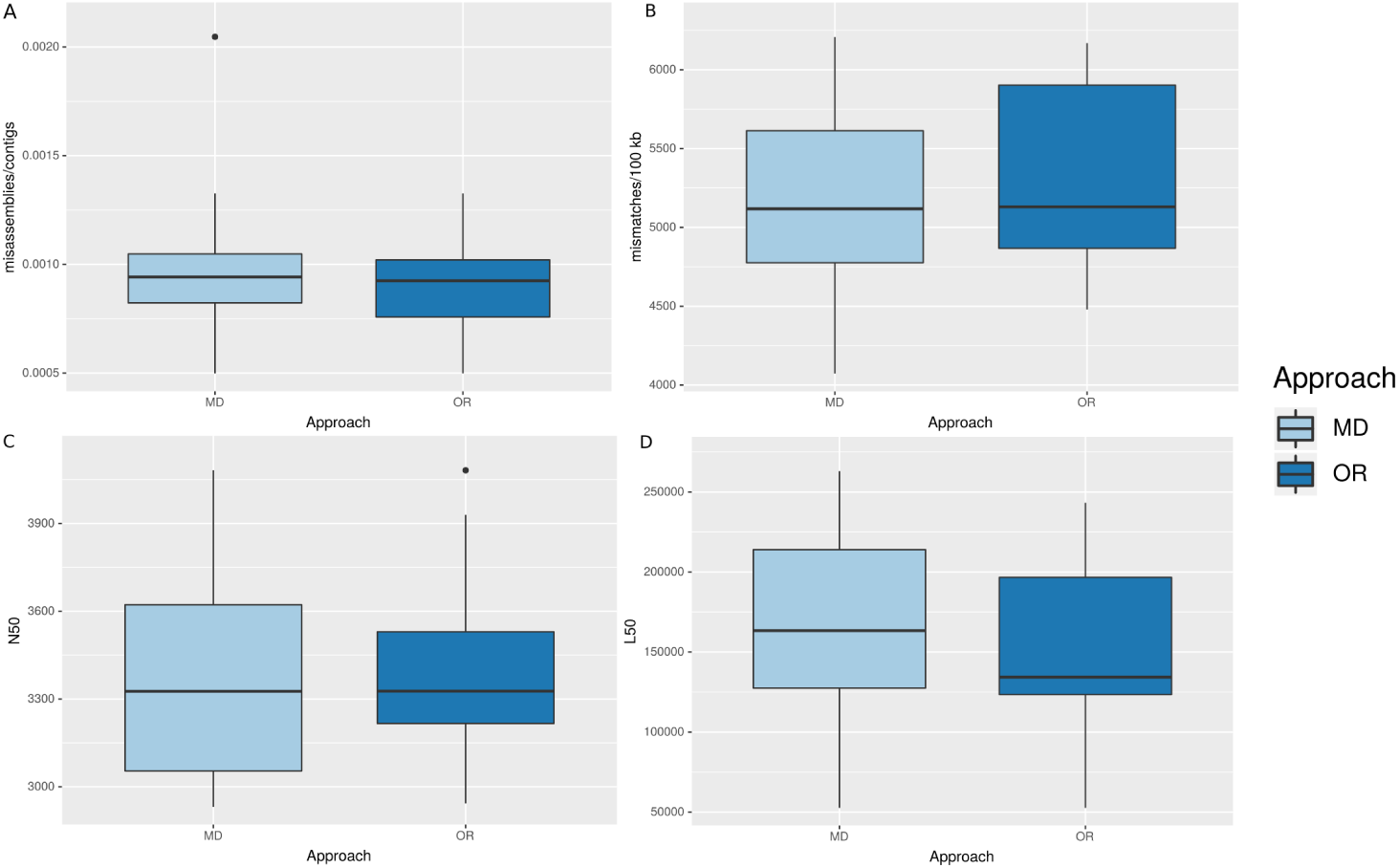
Quality of the assembly of each TARA approach. **A** Comparison of the number of misassemblies normalized by the number of contigs per assembly of the metagenomic distance (MD) and the oceanic regions (OR) approaches; **B** Number of mismatches detected per 100kb of alignments in contigs per approach; **C** N50 values: N50 represent the shortest contig length to cover at least 50% of the metagenome assembly; **D** L50 values: L50 represent the smallest number of contigs whose added lengths cover 50% of the metagenome assembly. No significant differences were found between the two approaches (Mann-Withney U test).

**FIG S3.**
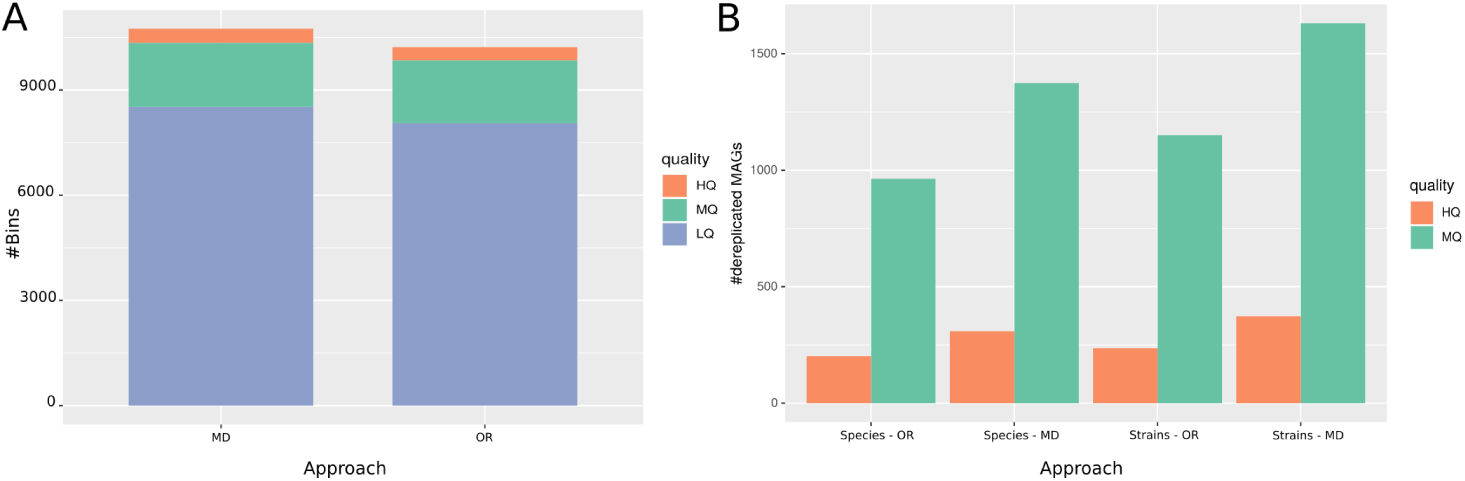
Evaluating the metagenomic distance-based (MD) approach against the Oceanic Regions (OR) approach for delineating groups of samples to co-assemble. **(A)** Total number of bins obtained after binning step. HQ = High Quality, MQ = Medium Quality, LQ = Low Quality. **(B)** Number of reconstructed MAGs after independent de-replication for each approach. OR = Oceanic Region, MD = Metagenomic Distance. Species resolution consists in a 95% ANI score de-replication, while Strains resolution consists in a 99% ANI score dereplication.

**FIG S4.**
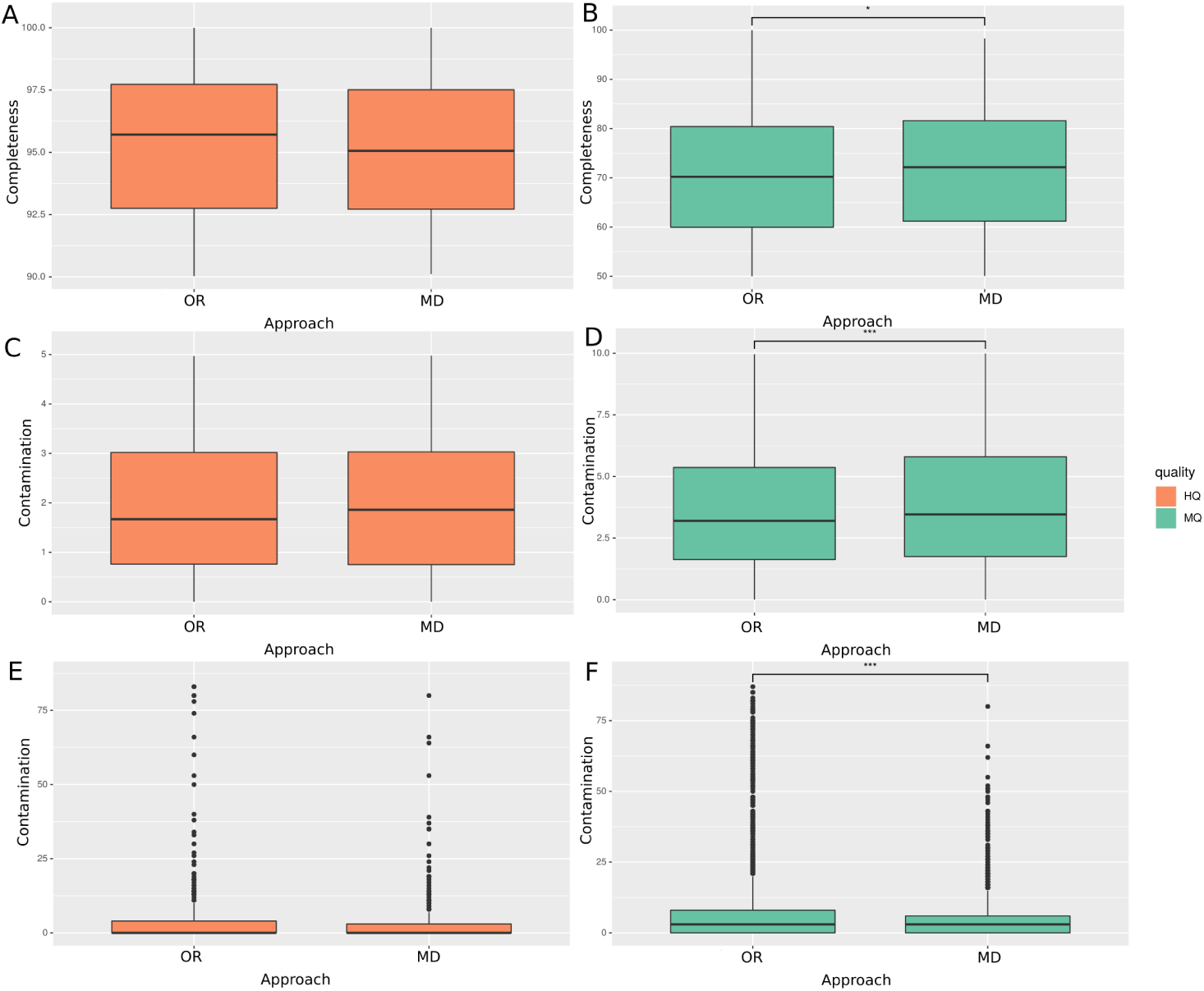
Quality of reconstructed MAGs from the marine metagenomes. Comparison of the oceanic region (OR) approach against the metagenomic distance approach (MD). HQ = High Quality, MQ = Medium Quality. **(A)** Completeness estimated for HQ MAGs; **(B)** Completeness estimated for MQ MAGs; **(C)** Contamination measured using SCGs for HQ MAGs; **(D)** Contamination measured using SCGs for MQ MAGs; **(E)** Contamination measured using all genes detected in the sequences of HQ MAGs; **(F)** Contamination measured using all genes detected in the sequences of MQ MAGs.

**FIG S5.**
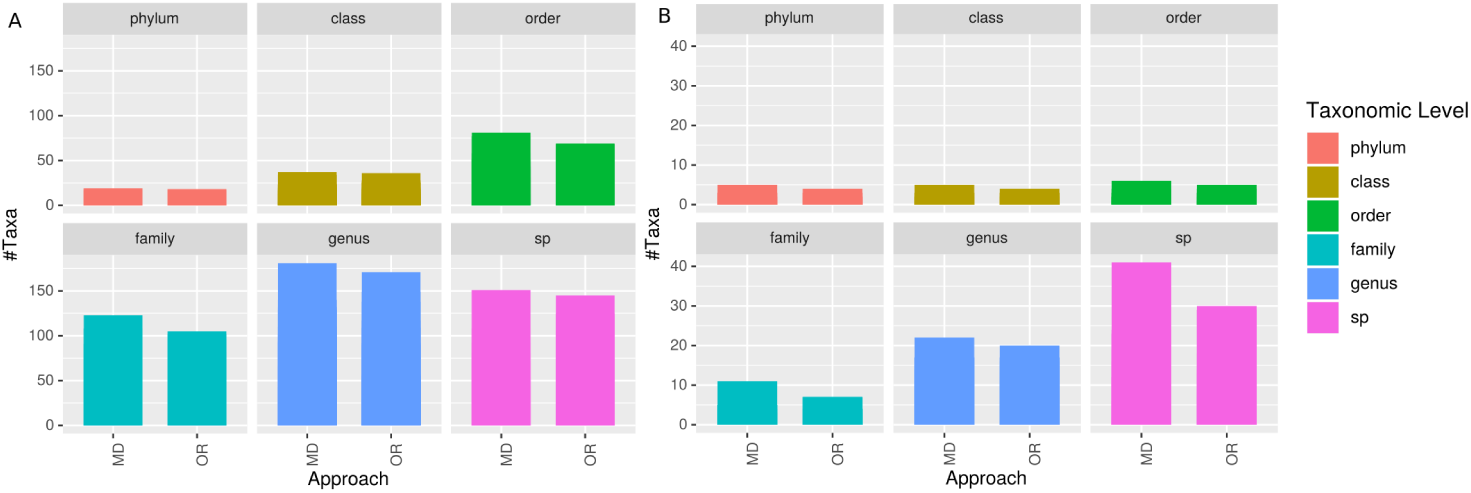
Taxonomic diversity of reconstructed MAGs from the marine metagenomes. Comparison of the number of taxa retrieved from MAGs reconstructed following each approach. **(A)**: Number of bacterial taxa assigned to MAGs from both approach, per taxonomic level ; **(B)**: Number of archaeal taxa assigned to MAGs from both approach. Taxonomic annotation was performed using gtdb-tk on de-replicated MAGs at strain level (99% ANI score) using dRep. For each panel, the y-axis represents the number of taxa found withing the complete set of MAGs of each approach. MD = Metagenomic Distance (estimated using Simka); OR = Oceanic Region (from (13)). Number of MAGs assigned to a bacterial annotation is 1729 for MD and 1595 for OR, while there are 276 and 273 MAGs with an archeal assignation in MD and OR, respectively.

**FIG S6.**
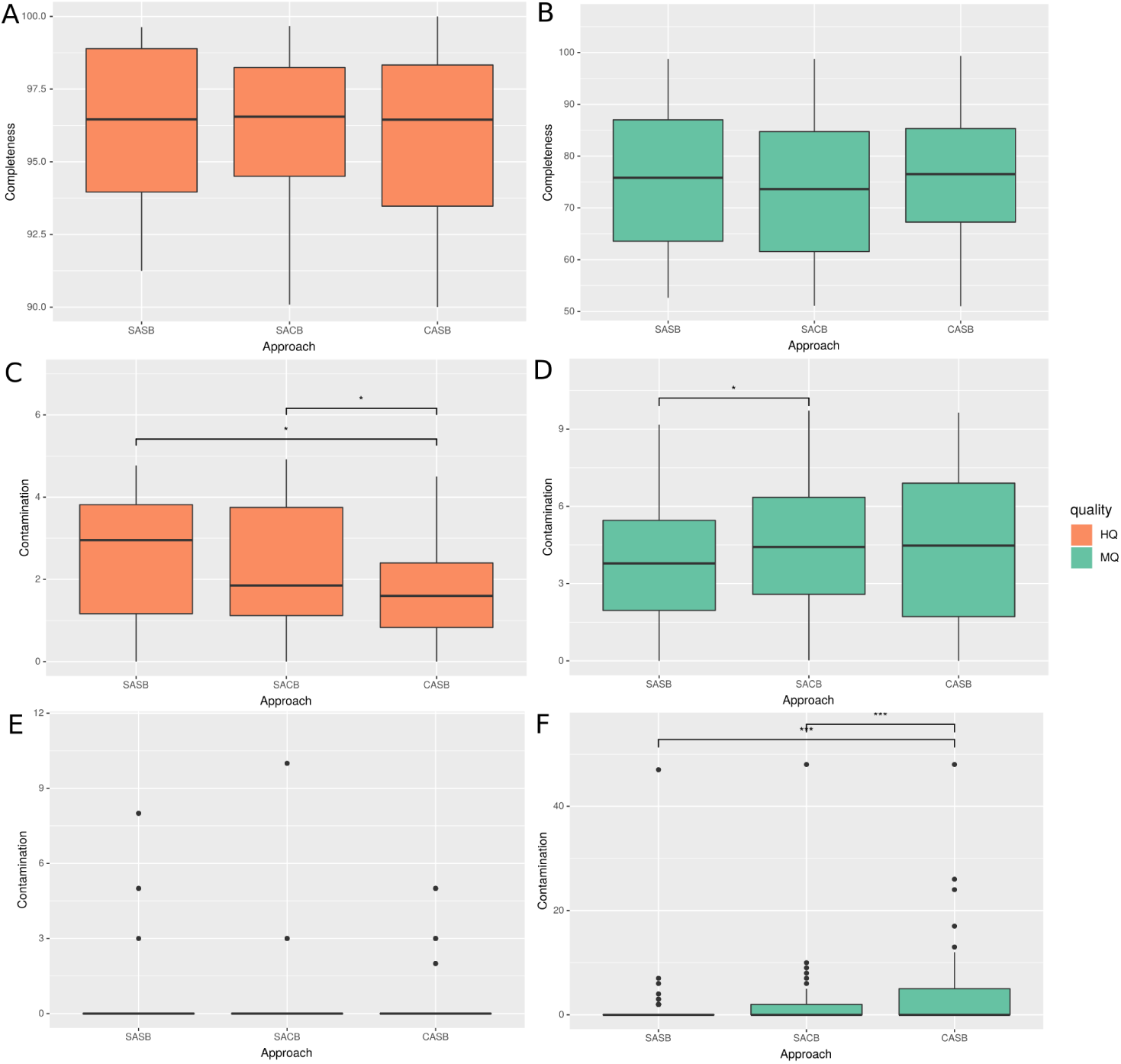
Quality of reconstructed MAGs from the CAMI dataset. Comparison of the quality of MAGs reconstructed with each strategy on the CAMI dataset. HQ = High Quality, MQ = Medium Quality. **(A)** Completeness estimated for HQ MAGs; **(B)** Completeness estimated for MQ MAGs; **(C)** Contamination measured using SCGs for HQ MAGs; **(D)** Contamination measured using SCGs for MQ MAGs; **(E)** Contamination measured using all genes detected in the sequences of HQ MAGs; **(F)** Contamination measured using all genes detected in the sequences of MQ MAGs.

**FIG S7.**
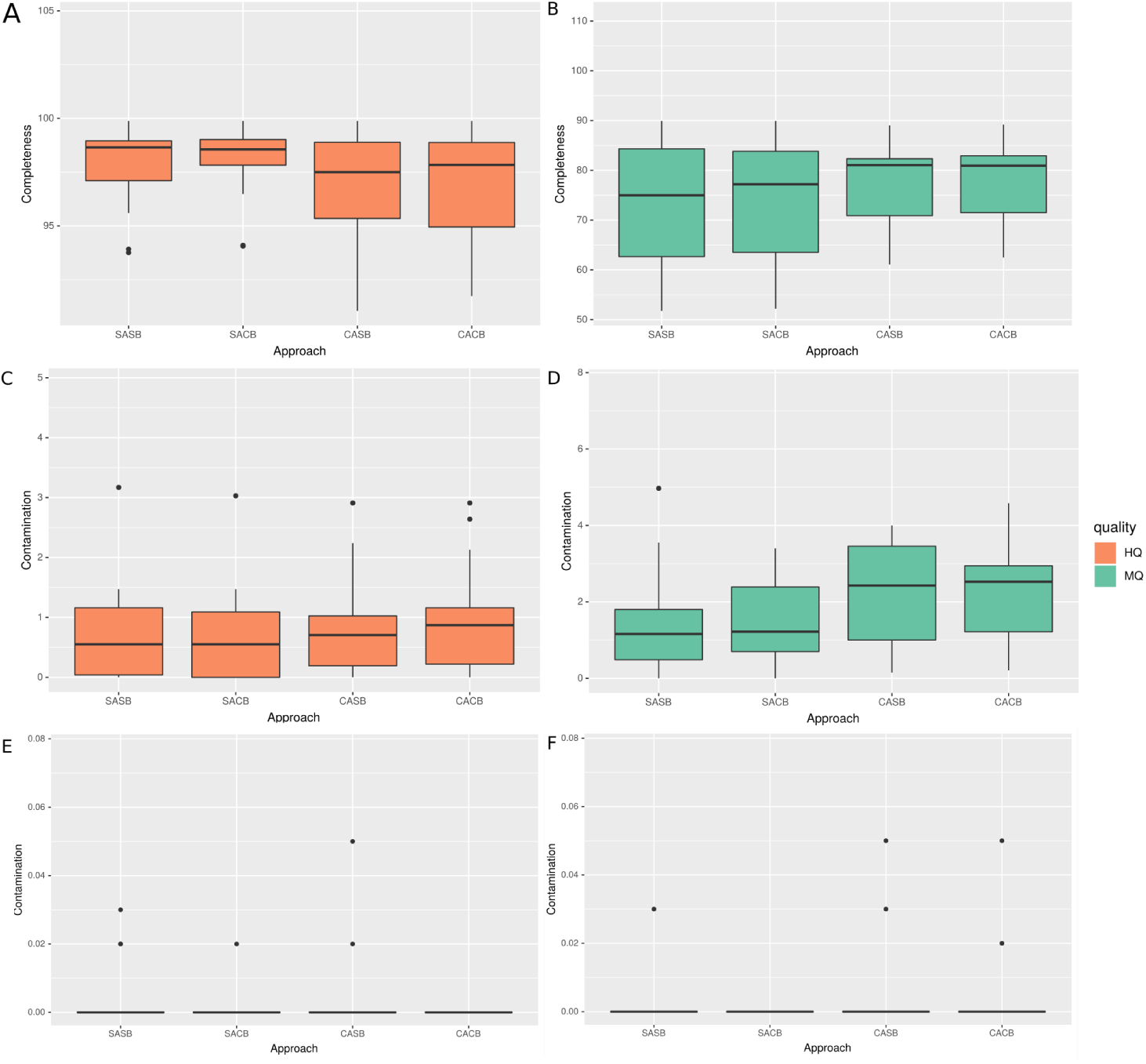
Quality of reconstructed MAGs from the mockdataset. Comparison of the quality of MAGs reconstructed with each strategy on the mockdataset. HQ = High Quality, MQ = Medium Quality. **(A)** Completeness of HQ MAGs; **(B)** Completeness of MQ MAGs; **(C)** Contamination measured with CheckM of HQ MAGs; **(D)** Contamination measured with CheckM of MQ MAGs, **(E)** Contamination of HQ MAGs, measured with GUNC; **(F)** Contamination of MQ MAGs, measured with GUNC.

**FIG S8.**
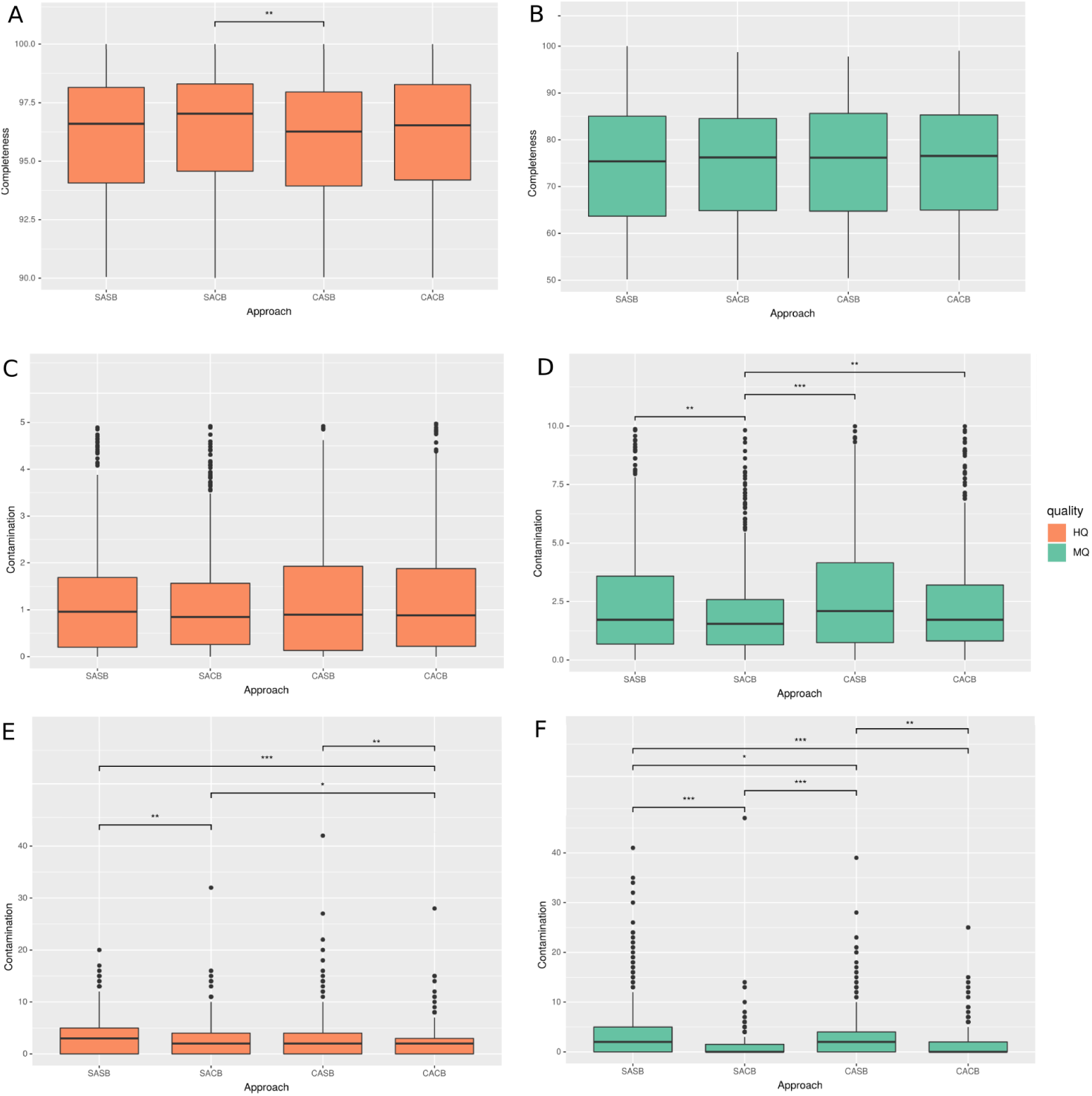
Quality of reconstructed MAGs from the HMP dataset. Comparison of the quality of MAGs reconstructed with each strategy on the HMP dataset. HQ = High Quality, MQ = Medium Quality. **(A)** Completeness estimated for HQ MAGs; **(B)** Completeness estimated for MQ MAGs; **(C)** Contamination measured using SCGs for HQ MAGs; **(D)** Contamination measured using SCGs for MQ MAGs; **(E)** Contamination measured using all genes detected in the sequences of HQ MAGs; **(F)** Contamination measured using all genes detected in the sequences of MQ MAGs.

**FIG S9.**
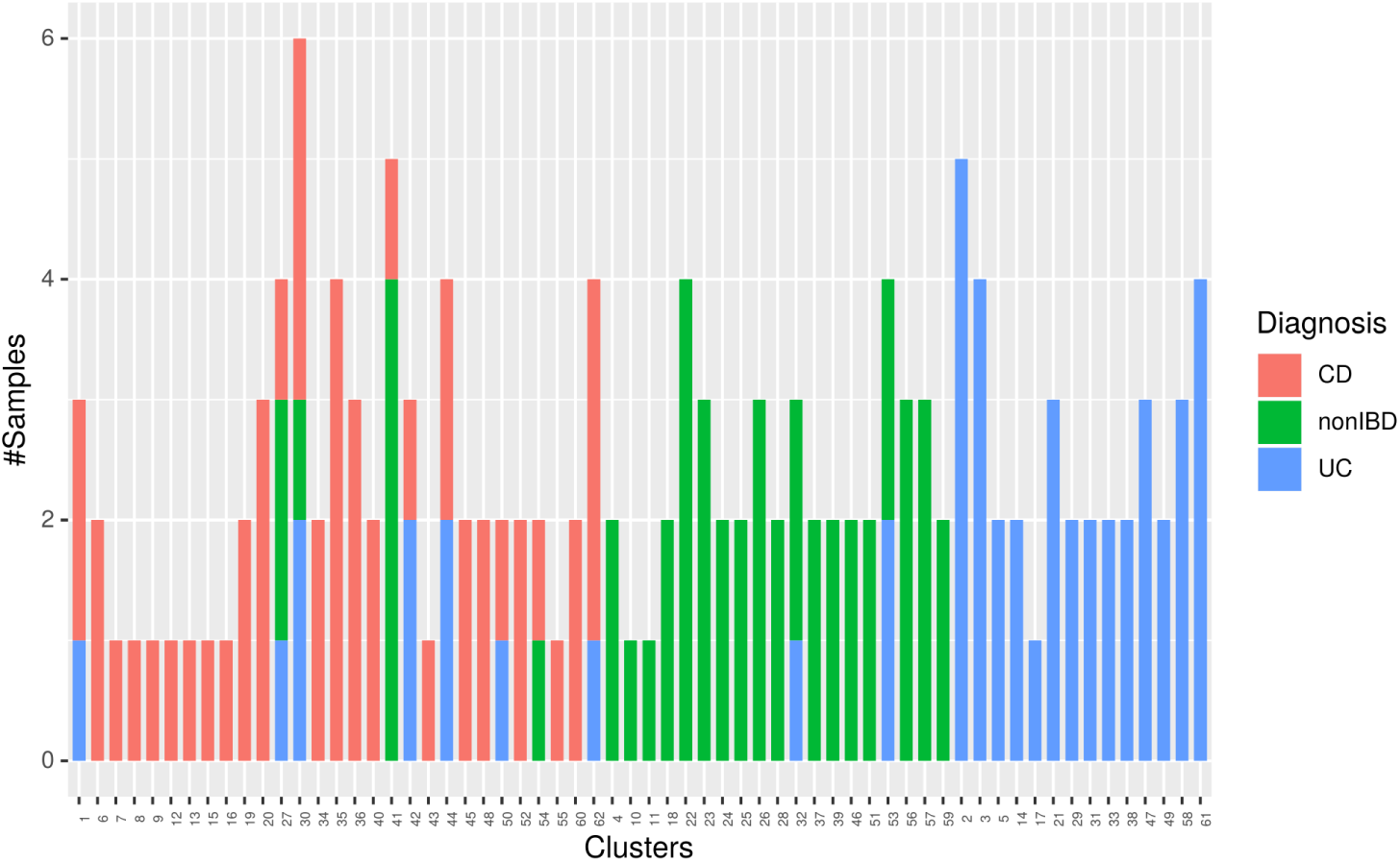
Composition of the MD clusters obtained with the HMP dataset. For each cluster, the amount of metagenomes is shown, with the IBD diagnosis associated with each metagenome. Diagnoses: non-IBD = healthy; CD = Crohn’s Disease; UC = Ulcerative Colitis

## REFERENCES

1. Kaeberlein T, Lewis K, Epstein SS. 2002. Isolating” uncultivable” microorganisms in pure culture in a simulated natural environment. Science 296 (5570):1127–1129.

2. Garza DR, Dutilh BE. 2015. From cultured to uncultured genome sequences: metagenomics and modeling microbial ecosystems. Cell Mol Life Sci 72 (22):4287–4308.

3. Almeida A, Nayfach S, Boland M, Strozzi F, Beracochea M, Shi ZJ, Pollard KS, Sakharova E, Parks DH, Hugenholtz P, et al. 2021. A unified catalog of 204,938 reference genomes from the human gut microbiome. Nat biotechnology 39 (1):105–114.

4. Hug LA, Baker BJ, Anantharaman K, Brown CT, Probst AJ, Castelle CJ, Butterfield CN, Hernsdorf AW, Amano Y, Ise K, et al. 2016. A new view of the tree of life. Nat microbiology 1 (5):1–6.

5. Nayfach S, Roux S, Seshadri R, Udwary D, Varghese N, Schulz F, Wu D, Paez-Espino D, Chen IM, Huntemann M, et al. 2021. Agenomic catalog of Earth’s microbiomes. Nat biotechnology 39 (4):499–509.

6. Albertsen M, Hugenholtz P, Skarshewski A, Nielsen KL, Tyson GW, Nielsen PH. Jun 2013. Genome sequences of rare, uncultured bacteria obtained by differential coverage binning of multiple metagenomes. Nat Biotechnol 31 (6):533–538. doi:10.1038/nbt.2579.

7. Alneberg J, Bjarnason BS, de Bruijn I, Schirmer M, Quick J, Ijaz UZ, Lahti L, Loman NJ, Andersson AF, Quince C. Nov 2014. Binning metagenomic contigs by coverage and composition. Nat Methods 11 (11):1144–1146. doi:10.1038/nmeth.3103.

8. Lu YY, Chen T, Fuhrman JA, Sun F. Jun 2016. COCACOLA: binning metagenomic contigs using sequence COmposition, read CoverAge, CO-alignment and paired-end read LinkAge. Bioinformatics pbtw290. doi:10.1093/bioinformatics/btw290.

9. Kang D, Li F, Kirton ES, Thomas A, Egan RS, An H, Wang Z. 2019. MetaBAT 2: an adaptive binning algorithm for robust and efficient genome reconstruction from metagenome assemblies. PeerJ Prepr 7:e27522v1.

10. Nelson WC, Tully BJ, Mobberley JM. 2020. Biases in genome reconstruction from metagenomic data. PeerJ 8:e10119.

11. Eren AM, Esen ÖC, Quince C, Vineis JH, Morrison HG, Sogin ML, Delmont TO. 2015. Anvi’o: an advanced analysis and visualization platform for ‘omics data. PeerJ 3:e1319.

12. Coleman I, Korem T. 2021. Embracing Metagenomic Complexity with a Genome-Free Approach. Msystems 6 (4):e00816–21.

13. Delmont TO, Quince C, Shaiber A, Esen OC, Lee ST, Rappé MS, McLellan SL, Lücker S, Eren AM. Jul 2018. Nitrogen-fixing populations of Planctomycetes and Proteobacteria are abundant in surface ocean metagenomes. Nat Microbiol 3 (7):804–813. doi: 10.1038/s41564-018-0176-9.

14. Pasolli E, Asnicar F, Manara S, Zolfo M, Karcher N, Armanini F, Beghini F, Manghi P, Tett A, Ghensi P, et al. 2019. Extensive unexplored human microbiome diversity revealed by over 150,000 genomes from metagenomes spanning age, geography, and lifestyle. Cell 176 (3):649–662.

15. Sczyrba A, Hofmann P, Belmann P, Koslicki D, Janssen S, Dröge J, Gregor I, Majda S, Fiedler J, Dahms E, et al. 2017. Critical as-sessment of metagenome interpretation—a benchmark of metagenomics software. Nat methods 14 (11):1063.

16. Wang Z, Wang Z, Lu YY, Sun F, Zhu S. 2019. SolidBin: improving metagenome binning with semi-supervised normalized cut. Bioinformatics 35 (21):4229–4238.

17. Herath D, Tang SL, Tandon K, Ackland D, Halgamuge SK. Dec 2017. CoMet: a workflow using contig coverage and composition for binning a metagenomic sample with high precision. BMC Bioinform 18 (S16). doi:10.1186/s12859-017-1967-3.

18. Graham ED, Heidelberg JF, Tully BJ. Mar 2017. BinSanity: unsupervised clustering of environmental microbial assemblies using coverage and affnity propagation. PeerJ 5:e3035. doi:10.7717/peerj.3035.

19. Nielsen HB, Almeida M, Juncker AS, Rasmussen S, Li J, Sunagawa S, Plichta DR, Gautier L, Pedersen AG, Le Chatelier E, et al. 2014. Identification and assembly of genomes and genetic elements in complex metagenomic samples without using reference genomes. Nat biotechnology 32 (8):822–828.

20. Plaza Oñate F, Le Chatelier E, Almeida M, Cervino AC, Gauthier F, Magoulès F, Ehrlich SD, Pichaud M. 2019. MSPminer: abundance-based reconstitution of microbial pan-genomes from shotgun metagenomic data. Bioinformatics 35 (9):1544–1552.

21. Mitchell AL, Almeida A, Beracochea M, Boland M, Burgin J, Cochrane G, Crusoe MR, Kale V, Potter SC, Richardson LJ, et al. 2020. MGnify: the microbiome analysis resource in 2020. Nucleic acids research 48 (D1):D570–D578.

22. Uritskiy GV, DiRuggiero J, Taylor J. Dec 2018. MetaWRAP—a flexible pipeline for genome-resolved metagenomic data analysis. Microbiome 6 (1). doi:10.1186/s40168-018-0541-1.

23. Clarke EL, Taylor LJ, Zhao C, Connell A, Lee JJ, Fett B, Bushman FD, Bittinger K. 2019. Sunbeam: an extensible pipeline for analyzing metagenomic sequencing experiments. Microbiome 7 (1):46.

24. Kieser S, Brown J, Zdobnov EM, Trajkovski M, McCue LA. 2020. ATLAS: a Snakemake workflow for assembly, annotation, and genomic binning of metagenome sequence data. BMC bioinformatics 21 (1):1–8.

25. Krakau S, Straub D, Gourlé H, Gabernet G, Nahnsen S. 2021. nfcore/mag: a best-practice pipeline for metagenome hybrid assembly and binning. bioRxiv doi:10.1101/2021.08.29.458094.

26. Paoli L, Ruscheweyh HJ, Forneris CC, Kautsar S, Clayssen Q, Salazar G, Milanese A, Gehrig D, Larralde M, Carroll LM, Sánchez P, Zayed AA, Cronin DR, Acinas SG, Bork P, Bowler C, Delmont TO, Sullivan MB, Wincker P, Zeller G, Robinson SL, Piel J, Sunagawa S. 2021. Uncharted biosynthetic potential of the ocean microbiome. bioRxiv doi:10.1101/2021.03.24.436479.

27. Clark DR, Underwood GJC, McGenity TJ, Dumbrell AJ. 2021. What drives study-dependent differences in distance–decay relationships of microbial communities? Glob Ecol Biogeogr 30 (4):811–825. doi: https://doi.org/10.1111/geb.13266.

28. Sunagawa S, Coelho LP, Chaffron S, Kultima JR, Labadie K, Salazar G, Djahanschiri B, Zeller G, Mende DR, Alberti A, Cornejo-Castillo FM, Costea PI, Cruaud C, d’Ovidio F, Engelen S, Ferrera I, Gasol JM, Guidi L, Hildebrand F, Kokoszka F, Lepoivre C, Lima-Mendez G, Poulain J, Poulos BT, Royo-Llonch M, Sarmento H, Vieira-Silva S, Dimier C, Picheral M, Searson S, Kandels-Lewis S, null null, Bowler C, de Vargas C, Gorsky G, Grimsley N, Hingamp P, Iudicone D, Jaillon O, Not F, Ogata H, Pesant S, Speich S, Stemmann L, Sullivan MB, Weissenbach J, Wincker P, Karsenti E, Raes J, Acinas SG, Bork P, Boss E, Bowler C, Follows M, Karp-Boss L, Krzic U, Reynaud EG, Sardet C, Sieracki M, Velayoudon D. 2015. Structure and function of the global ocean microbiome. Science 348 (6237):1261359. doi: 10.1126/science.1261359.

29. Richter DJ, Watteaux R, Vannier T, Leconte J, Frémont P, Reygondeau G, Maillet N, Henry N, Benoit G, Silva OD, Delmont TO, Fernàndez-Guerra A, Suweis S, Narci R, Berney C, Eveillard D, Gavory F, Guidi L, Labadie K, Mahieu E, Poulain J, Romac S, Roux S, Dimier C, Kandels S, Picheral M, Searson S, Coordinators TO, Pesant S, Aury JM, Brum JR, Lemaitre C, Pelletier E, Bork P, Sunagawa S, Lombard F, Karp-Boss L, Bowler C, Sullivan MB, Karsenti E, Mariadassou M, Probert I, Peterlongo P, Wincker P, de Vargas C, d’Alcalà MR, Iudicone D, Jaillon O. 2020. Genomic evidence for global ocean plankton biogeography shaped by large-scale current systems. bioRxiv doi:10.1101/867739.

30. Cabello-Yeves PJ, Callieri C, Picazo A, Mehrshad M, Haro-Moreno JM, Roda-Garcia JJ, Dzhembekova N, Slabakova V, Slabakova N, Moncheva S, et al. 2021. The microbiome of the Black Sea water column analyzed by shotgun and genome centric metagenomics. Environ microbiome 16 (1):1–15.

31. Karthikeyan S, Rodriguez-R LM, Heritier-Robbins P, Hatt JK, Huettel M, Kostka JE, Konstantinidis KT. 2020. Genome repository of oil systems: an interactive and searchable database that expands the catalogued diversity of crude oil-associated microbes. Environ Microbiol 22 (6):2094–2106.

32. Jégousse C, Vannier P, Groben R, Glöckner FO, Marteinsson V. 2021. A total of 219 metagenome-assembled genomes of microorganisms from Icelandic marine waters. PeerJ 9:e11112.

33. Vosloo S, Huo L, Anderson CL, Dai Z, Sevillano M, Pinto A. 2021. Evaluating de Novo Assembly and Binning Strategies for Time Series Drinking Water Metagenomes. Microbiol spectrum 9 (3):e01434–21.

34. Merrill BD, Carter MM, Olm MR, Dahan D, Tripathi S, Spencer SP, Feiqiao BY, Jain S, Neff N, Jha AR, et al. 2022. Ultra-deep Sequencing of Hadza Hunter-Gatherers Recovers Vanishing Microbes. bioRxiv.

35. Benoit G, Peterlongo P, Mariadassou M, Drezen E, Schbath S, Lavenier D, Lemaitre C. Nov 2016. Multiple comparative metagenomics using multiset k -mer counting. PeerJ Comput Sci 2:e94. doi: 10.7717/peerj-cs.94.

36. Rousseeuw PJ. 1987. Silhouettes: a graphical aid to the interpretation and validation of cluster analysis. J computational applied mathematics 20:53–65.

37. Olm MR, Brown CT, Brooks B, Banfield JF. 2017. dRep: a tool for fast and accurate genomic comparisons that enables improved genome recovery from metagenomes through de-replication. The ISME journal 11 (12):2864.

38. Orakov A, Fullam A, Coelho LP, Khedkar S, Szklarczyk D, Mende DR, Schmidt TS, Bork P. 2021. GUNC: detection of chimerism and contamination in prokaryotic genomes. Genome biology 22 (1):1–19.

39. Fritz A, Hofmann P, Majda S, Dahms E, Dröge J, Fiedler J, Lesker TR, Belmann P, DeMaere MZ, Darling AE, et al. 2019. CAMISIM: simulating metagenomes and microbial communities. Microbiome 7 (1):1–12.

40. Lloyd-Price J, Arze C, Ananthakrishnan AN, Schirmer M, Avila-Pacheco J, Poon TW, Andrews E, Ajami NJ, Bonham KS, Brislawn CJ, et al. 2019. Multi-omics of the gut microbial ecosystem in inflammatory bowel diseases. Nature 569 (7758):655–662.

41. Li D, Liu CM, Luo R, Sadakane K, Lam TW. 2015. MEGAHIT: an ultrafast single-node solution for large and complex metagenomics assembly via succinct de Bruijn graph. Bioinformatics 31 (10):1674–1676.

42. Sieber CMK, Probst AJ, Sharrar A, Thomas BC, Hess M, Tringe SG, Banfield JF. Jul 2018. Recovery of genomes from metagenomes via a dereplication, aggregation and scoring strategy. Nat Microbiol 3 (7):836–843. doi:10.1038/s41564-018-0171-1.

43. Köster J, Rahmann S. 2012. Snakemake—a scalable bioinformatics workflow engine. Bioinformatics 28 (19):2520–2522.

44. Chen S, Zhou Y, Chen Y, Gu J. 09 2018. fastp: an ultra-fast all-in-one FASTQ preprocessor. Bioinformatics 34 (17):i884–i890. doi: 10.1093/bioinformatics/bty560.

45. Wingett SW, Andrews S. 2018. FastQ Screen: A tool for multigenome mapping and quality control. F1000Research 7.

46. Parks DH, Imelfort M, Skennerton CT, Hugenholtz P, Tyson GW. 2015. CheckM: assessing the quality of microbial genomes recovered from isolates, single cells, and metagenomes. Genome research 25 (7):1043–1055.

47. Steinegger M, Söding J. Dec 2018. Clustering huge protein sequence sets in linear time. Nat Commun 9 (1). doi:10.1038/s41467-018-04964-5.

48. Chaumeil PA, Mussig AJ, Hugenholtz P, Parks DH. 2020. GTDB-tk: a toolkit to classify genomes with the Genome Taxonomy Database. Oxford University Press.

49. Huerta-Cepas J, Szklarczyk D, Heller D, Hernández-Plaza A, Forslund SK, Cook H, Mende DR, Letunic I, Rattei T, Jensen LJ, et al. 2019. eggNOG 5.0: a hierarchical, functionally and phylogenetically annotated orthology resource based on 5090 organisms and 2502 viruses. Nucleic acids research 47 (D1):D309–D314.

50. Logares R, Deutschmann IM, Junger PC, Giner CR, Krabberød AK, Schmidt TS, Rubinat-Ripoll L, Mestre M, Salazar G, Ruiz-González C, et al. 2020. Disentangling the mechanisms shaping the surface ocean microbiota. Microbiome 8 (1):1–17.

51. Kerkhof L, Voytek M, Sherrell RM, Millie D, Schofield O. 1999. Variability in bacterial community structure during upwelling in the coastal ocean. Hydrobiologia 401:139–148.

52. Allen LZ, Allen EE, Badger JH, McCrow JP, Paulsen IT, Elbourne LD, Thiagarajan M, Rusch DB, Nealson KH, Williamson SJ, et al. 2012. Influence of nutrients and currents on the genomic composition of microbes across an upwelling mosaic. The ISME journal 6 (7):1403–1414.

53. Charuvaka A, Rangwala H. 2011. Evaluation of short read metagenomic assembly. BMC Genom 12 (2):S8.

54. Le Gall G, Noor SO, Ridgway K, Scovell L, Jamieson C, Johnson IT, Colquhoun IJ, Kemsley EK, Narbad A. 2011. Metabolomics of fecal extracts detects altered metabolic activity of gut microbiota in ulcerative colitis and irritable bowel syndrome. J proteome research 10 (9):4208–4218.

55. Zheng D, Liwinski T, Elinav E. 2020. Interaction between microbiota and immunity in health and disease. Cell research 30 (6):492–506.

56. Flores GE, Caporaso JG, Henley JB, Rideout JR, Domogala D, Chase J, Leff JW, Vázquez-Baeza Y, Gonzalez A, Knight R, et al. 2014. Temporal variability is a personalized feature of the human microbiome. Genome biology 15 (12):1–13.

57. Gilbert JA, Blaser MJ, Caporaso JG, Jansson JK, Lynch SV, Knight R. 2018. Current understanding of the human microbiome. Nat medicine 24 (4):392–400.

58. Sangwan N, Xia F, Gilbert JA. Dec 2016. Recovering complete and draft population genomes from metagenome datasets. Microbiome 4 (1). doi:10.1186/s40168-016-0154-5.

59. Nurk S, Meleshko D, Korobeynikov A, Pevzner PA. 2017. metaSPAdes: a new versatile metagenomic assembler. Genome research 27 (5):824–834.

60. Forouzan E, Shariati P, Maleki MSM, Karkhane AA, Yakhchali B. 2018. Practical evaluation of 11 de novo assemblers in metagenome assembly. J microbiological methods 151:99–105.

61. Vollmers J, Wiegand S, Kaster AK. 2017. Comparing and evaluating metagenome assembly tools from a microbiologist’s perspective-not only size matters! PloS one 12 (1):e0169662.

62. Ward Jr JH. 1963. Hierarchical grouping to optimize an objective function. J Am statistical association 58 (301):236–244.

63. Langmead B, Salzberg SL. 2012. Fast gapped-read alignment with Bowtie 2. Nat methods 9 (4):357.

64. Kang DD, Froula J, Egan R, Wang Z. Aug 2015. MetaBAT, an efficient tool for accurately reconstructing single genomes from complex microbial communities. PeerJ 3:e1165. doi:10.7717/peerj.1165.

65. Bowers RM, Kyrpides NC, Stepanauskas R, Harmon-Smith M, Doud D, Reddy TBK, Schulz F, Jarett J, Rivers AR, Eloe-Fadrosh EA, Tringe SG, Ivanova NN, Copeland A, Clum A, Becraft ED, Malmstrom RR, Birren B, Podar M, Bork P, Weinstock GM, Garrity GM, Dodsworth JA, Yooseph S, Sutton G, Glöckner FO, Gilbert JA, Nelson WC, Hallam SJ, Jungbluth SP, Ettema TJG, Tighe S, Konstantinidis KT, Liu WT, Baker BJ, Rattei T, Eisen JA, Hedlund B, McMahon KD, Fierer N, Knight R, Finn R, Cochrane G, Karsch-Mizrachi I, Tyson GW, Rinke C, Kyrpides NC, Schriml L, Garrity GM, Hugenholtz P, Sutton G, Yilmaz P, Meyer F, Glöckner FO, Gilbert JA, Knight R, Finn R, Cochrane G, Karsch-Mizrachi I, Lapidus A, Meyer F, Yilmaz P, Parks DH, Eren AM, Schriml L, Banfield JF, Hugenholtz P, Woyke T. Aug 2017. Minimum information about a single amplified genome (MISAG) and a metagenome-assembled genome (MIMAG) of bacteria and archaea. Nat Biotechnol 35 (8):725–731. doi: 10.1038/nbt.3893.

66. R Core Team. 2018. R: A Language and Environment for Statistical Computing. R Foundation for Statistical Computing, Vienna, Austria. https://www.R-project.org/.

67. Varghese NJ, Mukherjee S, Ivanova N, Konstantinidis KT, Mavrommatis K, Kyrpides NC, Pati A. 2015. Microbial species delineation using whole genome sequences. Nucleic acids research 43 (14):6761–6771.

68. Olm MR, Crits-Christoph A, Diamond S, Lavy A, Matheus Carnevali PB, Banfield JF. 2020. Consistent metagenome-derived metrics verify and delineate bacterial species boundaries. Msystems 5 (1):e00731–19.

69. Hyatt D, Chen GL, LoCascio PF, Land ML, Larimer FW, Hauser LJ. 2010. Prodigal: prokaryotic gene recognition and translation initiation site identification. BMC bioinformatics 11 (1):119.

70. Steinegger M, Söding J. 2017. MMseqs2 enables sensitive protein sequence searching for the analysis of massive data sets. Nat biotechnology 35 (11):1026–1028.

